# Eco-Evolutionary Consequences of Selective Exploitation on Metapopulations Illustrated With Atlantic Salmon

**DOI:** 10.1101/2024.07.25.605173

**Authors:** Amaïa Lamarins, Stephanie M. Carlson, Etienne Prévost, William H. Satterthwaite, Mathieu Buoro

## Abstract

While the eco-evolutionary consequences of dispersal and exploitation are increasingly recognized, consideration of these effects and how they interact for management and conservation remains limited. We addressed this gap by examining population exploitation within a metapopulation framework, using Atlantic salmon as a case study. We compared eco-evolutionary consequences of alternative exploitation strategies by incorporating selective exploitation based on life history traits and spatial dimension of exploitation (i.e., whether populations were net exporter or importer of individuals). We used a demo-genetic agent-based model to examine demographic and evolutionary consequences of these strategies across a gradient of population-specific exploitation rates. At the metapopulation scale, we found both lower abundance and earlier sexual maturation with increasing exploitation, particularly when fishing was selective on larger individuals. The spatial selectivity of exploitation had an overall additional detrimental effect on metapopulation performance and fisheries yield, and induced stronger evolutionary changes than when exploitation was evenly spread over all populations. We discuss the implications of metapopulation functioning for species management and how considering dispersal patterns and intensity might change how we apply harvest. Nevertheless, our findings suggest that the safest approach remains to distribute exploitation efforts evenly across all populations, especially in the absence of variation in intrinsic productivity and with the dispersal rates and spatial configuration simulated. However, this strategy might not completely prevent negative consequences at the local scale. Therefore, we advise managers to critically assess the relevance of our results and dispersal assumptions in the specific cases they may have to deal with.

## 1 Introduction

Natural resources exploitation and conservation are increasingly at odds in the context of global change, prompting management authorities and scientists to reconsider how to exploit species without threatening their long-term persistence. Management strategies should aim to ensure the sustainability of stock and yield, while also promoting resilience and adaptation of the exploited populations in a changing environment. It is thus important to not only consider the demographic status of populations, but also to integrate an evolutionary perspective in management decisions (Hoffmann et al., 2015), by explicitly considering the eco-evolutionary consequences of exploitation.

Beyond its direct demographic effects, exploitation can act as a selective force (Fenberg and Roy, 2008). In fisheries, there is often a preference to harvest larger individuals, which can negatively affect productivity and diversity of the targeted populations (Garcia et al., 2012). It can also cause evolutionary changes (Olsen et al., 2004; Allendorf and Hard, 2009; Czorlich et al., 2022) and reduce genetic diversity (Smith et al., 1991). This phenomenon is termed “fisheries-induced evolution” (Kuparinen and Merilä, 2007; Heino et al., 2015) and its potential consequences for the demographic recovery of fish stocks is well-documented (Enberg et al., 2009). Ultimately, the selective exploitation of a population can affect its eco-evolutionary dynamic, because the evolutionary changes can feed back into ecological processes and affect demography (Govaert et al., 2019).

It is also possible for mortality induced by exploitation to be selective not only *within* populations but also *across* populations. The exploitation of a set of conspecific populations has been extensively studied in the case of mixed-stock fisheries that jointly exploit as a “single” large stock several demographically-distinct populations gathered in the same location (Ricker, 1958; Hilborn et al., 2003). In a mixed-stock fishery, exploitation is usually distributed among the populations in ways that may not be proportionate to size or risk status. It may lead to overexploitation of the weaker stocks, thus increasing their risk of collapse. The dangers of ignoring the status of individual populations have been highlighted in numerous studies, e.g., on cod (Hutchinson, 2008), sockeye salmon (Moore et al., 2021), and croaker (Ying et al., 2011). Managing fisheries in a way that considers the characteristics and vulnerabilities of individual stocks is most difficult when they are mixed and fishing takes place in locations of mixing. But when populations are both demographically distinct and spatially segregated, for example in various rivers or lakes or at the time of river return for anadromous fish, there is greater opportunity to manage exploitation at the level of local populations.

However, even when conspecific populations are spatially segregated, they are rarely isolated from their neighbours as they are often connected by movements of individuals through dispersal (Clobert et al., 2001, 2012). This results in spatially structured populations and metapopulation functioning which is critical to consider in species conservation efforts (Akçakaya et al., 2007). Focusing solely on the impact of exploitation on a single population may be misleading, as local abundance does not always reflect the local productivity due to substantial net emigration or immigration from other populations. Additionally, local exploitation of the most productive populations can affect interconnected populations, by reducing demographic subsidies to other populations. Dispersal plays an essential role in eco-evolutionary dynamics of populations because it allows demographic, genetic and evolutionary rescue effects (Brown and Kodric-Brown, 1977; Carlson et al., 2014). Thus, it could dampen the evolutionary and demographic consequences of selective exploitation, contributing to metapopulation stability. Moreover, dispersal can lead to gene flow, which can alter local adaptation and traits connected to yield (e.g., life history tactic), suggesting multiple reasons for better understanding consequences of selective exploitation in a metapopulation context.

There is still limited knowledge about the most appropriate management approaches to adopt for a set of interconnected populations with conflicting objectives of conservation (e.g., population persistence and genetic diversity) and exploitation. Explicitly considering metapopulation dynamics, some studies advocate for the protection of certain populations over others based on various criteria such as population size (Okamoto et al., 2020), whether they are a source or a sink (Crowder et al., 2000; Tufto and Hindar, 2003; Fullerton et al., 2016), their influence on spatial synchrony (Engen et al., 2018), or connectivity metrics (Kininmonth et al., 2019; Radici et al., 2023). Nevertheless, despite the widespread recognition of selective exploitation in fisheries, the literature has not addressed the eco-evolutionary impact of selective exploitation of fitness-related traits within a framework of evolving metapopulations. Developing harvest rules for spatially complex fish populations remains a challenge (Benson et al., 2015), and the explicit consideration of evolutionary processes, together with metapopulation functioning, may provide new insights into the sustainable exploitation of metapopulations.

However, investigating the eco-evolutionary consequences of selective exploitation and management strategies in wild metapopulations by real-world experiments or observational studies can be challenging, if not impossible. Simulation models provide unrivalled opportunities to assess and compare the performance of alternative management actions and anticipate poor outcomes without risking invaluable resources. Different modeling approaches can be used (Pelletier and Mahévas, 2005; Zurell et al., 2022), each with benefits and limitations depending on the management context. But most of the spatially explicit simulation models published so far in fisheries science ignore genetics and evolutionary processes (e.g., Okamoto et al., 2020), limiting our understanding of the impacts of selective exploitation on populations. Demo-genetic Agent Based Models (DG-ABMs), defined as “individual-based (meta) population dynamics models with heritable trait variation and phenotype-dependent interactions between individuals” (Lamarins et al., 2022*a*), have emerged as an effective framework to study management strategies while being spatially explicit and integrating eco-evolutionary feedbacks. DG-ABMs have been used to investigate the consequences of exploitation but mainly at population scale (e.g., Thériault et al., 2008; Marty et al., 2015; Ayllón et al., 2019; Piou et al., 2015). Recent spatially explicit DG-ABMs such as MetaIBASAM, a model that simulates spatially structured Atlantic salmon populations, have opened up new possibilities (Lamarins et al., 2022*b*).

Identifying harvest management strategies that achieve conservation goals in the context of evolving metapopulations submitted to selective fisheries is of particular interest for salmonids. It is especially true for Atlantic salmon (*Salmo salar*), a species harvested for centuries (Aas et al., 2010) by a diversity of indigenous, commercial and recreational fisheries operating in rivers, estuaries and at sea, and emblematic of the tension between conservation and exploitation. Its overall abundance has declined drastically throughout its area of distribution over the last decades (ICES, 2021), mainly due to alteration of within-river connectivity hampering its migrations, reduced marine survival, and overexploitation (Chaput, 2012), the latter reaching rates up to 80% in the West Greenland fishery in the 1980s (Olmos et al., 2019). In an attempt to halt the decline of Atlantic salmon, the main commercial fisheries at sea were closed in the 1990s and now most of the exploitation occurs in rivers and estuaries (terminal fisheries, ICES, 2021). Atlantic salmon is an anadromous and philopatric species that divides its life cycle between freshwater for egg incubation and juvenile development and the ocean for growth before returning, for the most part, to its natal river to spawn. Consequently, its populations are often managed as distinct and independent between rivers, with exploitation rates that often vary among nearby populations (e.g., Dempson et al., 2001, Bradbury et al., 2015). It is known, however, that some individuals disperse to and reproduce in other rivers than their natal one (Jonsson et al., 2003; Consuegra et al., 2005), and earlier studies have warned against the “danger of ignoring metapopulation structure for conservation” of salmonids (Cooper and Mangel, 1999; Schtickzelle and Quinn, 2007; Bradford and Braun, 2021). At the same time, there is an overall tendency of Atlantic salmon fisheries to be selective among returning adults according to their life histories, in particular maturation age. Fish may mature after one winter at sea (1SW), or after multiple winters at sea (MSW), and higher exploitation rates have been often reported for the latter (e.g., Dempson et al., 2001; Thorley et al., 2007; Lebot et al., 2022).

The identification of harvest management strategies that achieve conservation goals in the context of evolving meta-populations is undoubtedly useful and needed. Our study aims to elucidate the demographic and evolutionary consequences of selective exploitation in spatially structured Atlantic salmon populations using a simulation approach. We investigate alternative strategies of exploitation that are i) selective or non-selective on a life history trait related to fitness (maturation age) and ii) spatially selective or non-selective on populations based on their source / sink status, defined from net migration (Pulliam, 1988; Loreau et al., 2013) and independent from variation in intrinsic productivity. We use MetaIBASAM (Lamarins et al., 2022*b*), a DG-ABM to simulate Atlantic salmon metapopulations exposed to various intensities of selective exploitation on a life-history trait (*within* population) and in space (*across* source-sink populations). We first assess the consequences of life history selectivity by comparing strategies increasingly selective against the older and larger (MSW) individuals with strategies that are non-selective. We then explore strategies of spatial selectivity, by exploiting populations based on whether they are source populations, sink populations, or neutral (asymmetry in net migration induced by differences in habitat size and spatial arrangement), in order to evaluate the effects of population type in a life history non-selective vs. selective exploitation context. We examine the demographic and evolutionary consequences of these strategies and contrast their performance relative to both metapopulation performance (i.e., abundance, stability, extinction risk) and exploitation goals. Selective exploitation of traits within a population is expected to induce evolutionary changes and exacerbate metapopulation reduction compared to non-selective exploitation, due to eco-evolutionary feedbacks. We also expect spatial selectivity of exploitation to affect the dynamics of the metapopulation, its resilience and stability, by limiting or favouring rescue effects.

## 2 Methods

### 2.1 A spatially explicit demo-genetic agent-based model

Piou and Prévost (2012) designed a DG-ABM to assess the effects of environmental change and selective fisheries on the demography and life history traits of a single population of Atlantic salmon (IBASAM, Piou and Prévost, 2013; Piou et al., 2015). It was further extended by (2022*b*) to a metapopulation framework (MetaIBASAM), which simulates the eco-evolutionary dynamics of interconnected populations. The MetaIBASAM model incorporates both demographic and genetic processes, as well as their interactions; represents explicitly each individual with its traits (e.g., size, growth potential), and potential interactions with others (e.g., density-dependence, sexual selection); and considers the populations’ spatial configuration and heterogeneity. Below we provide a brief overview of the model structure and additional details are available in the previously cited papers.

For each population within the network, the life cycle of each individual – which encapsulates the eco-evolutionary processes such as emergence, growth, survival, migration, maturation and reproduction – is simulated with a daily time step. The biological traits (e.g., sexual maturity, age, size) of each individual are monitored. Atlantic salmon is an anadromous species with a great diversity of life history strategies (Erkinaro et al., 2019). Reproduction and juvenile growth occur in rivers, and in the southern part of their distribution area, two successive life history decisions are made: to mature sexually in freshwater or not (precocious maturation occurring only in “parr” males to reproduce in the upcoming spawning season), and to migrate to the sea after their first or second year of life (a physiological transformation to a “smolt” stage). At sea, individuals grow and make an additional life history decision: to mature after only one year at sea (“one sea-winter” or 1SW) or multiple years (“multiple sea-winter” or MSW) before returning to rivers to breed as anadromous adults. A high proportion of maturing adults return to their river of birth (philopatry), but dispersal to non-natal rivers also occurs regularly (Jonsson et al., 2003; Consuegra et al., 2005). A proportion of adults returning to each river is removed by fishing (terminal fisheries in freshwater) and in the model, the exploitation rate (i.e., probability of capture) depends only on the river of return (i.e., after immigration), the time spent at sea, and the year. Exploitation can thus be set up as selective according to sea age category (Piou et al., 2015) and according to space by targeting populations with specific characteristics.

Environmental forcing (e.g., water temperature, water flow) and density-dependent processes affect life history and demographic processes, mainly river growth and survival (Suppl. Mat. Fig. S1). A set of traits (growth potential, maturation thresholds triggering sexual maturation in river and at sea) also include an underlying genetic basis, allowing evolution of traits in response to selection, and emergence of eco-evolutionary feedbacks. The growth potential represents the individual capacity to grow and may be under selection; for example, the decision to migrate to the sea is based on a probabilistic size-dependent reaction norm, while the survival at sea is also size-dependent. Maturation decision in river (for male “parr”) or at sea (for anadromous adults) is based on an environmental threshold model, i.e., the maturation occurs when an individual’s energetic reserves (based on growth) exceed its maturation threshold (genetically based). Local populations are connected by dispersal of individuals, which occurs when adults are migrating from the sea to the rivers, before fishing and reproduction. Dispersal patterns are defined via a fixed emigration rate (i.e., the probability of returning to a non-natal river) and a dispersal kernel determining the recipient populations according to their distance from the natal river and their carrying capacity (i.e., a measure of “attractivity”).

The model was parameterized in a pattern-oriented modeling framework (Grimm et al., 2005) as in Piou and Prévost (2012) and Lamarins et al. (2022*b*), and drew from the spatial structure 219 (i.e., distance and carrying capacities) of the salmon metapopulation of Brittany (France, Perrier et al., 2011; Bouchard et al., 2022), i.e. small coastal rivers. The network spatial configuration was constituted of fifteen adjacent populations, i.e. not following a dendritic network (Fig. 1).

**Figure 1:**
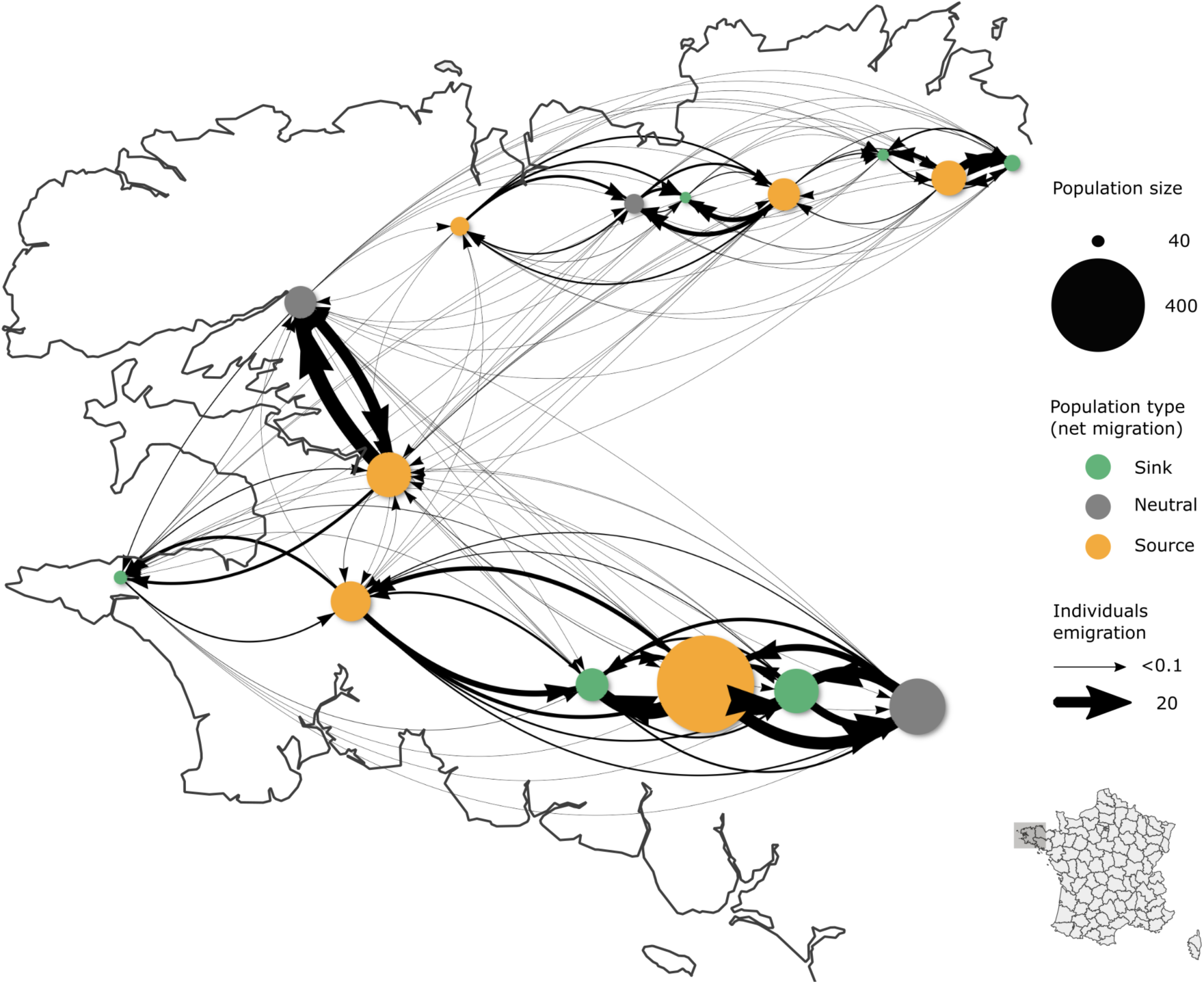
Spatial configuration of the network used in this analysis based on the Brittany metapopulation. Populations are represented by circles (size scaled to population size, color based on net migration) and emigration flows between populations are represented by the arrows (width scaled to emigrants number). Modified from Lamarins et al. (2022*b*) with a dispersal rate of 15% and no exploitation.

### 2.2 Scenarios of exploitation

We focus on the consequences of different strategies of exploitation, i) selective vs non-selective exploitation based on life history traits (1SW vs MSW), and ii) spatial selectivity of exploitation based on population types (Fig. 2). For non-selective exploitation on life histories, we tested scenarios following a gradient of local exploitation rates from 0 to 100%, irrespective of adult traits. For selective exploitation on life histories, we tested scenarios maintaining a local exploitation rate of 10% on 1SW individuals and following a gradient of local exploitation rates from 0 to 100% on MSW individuals (in 10% increments).

**Figure 2:**
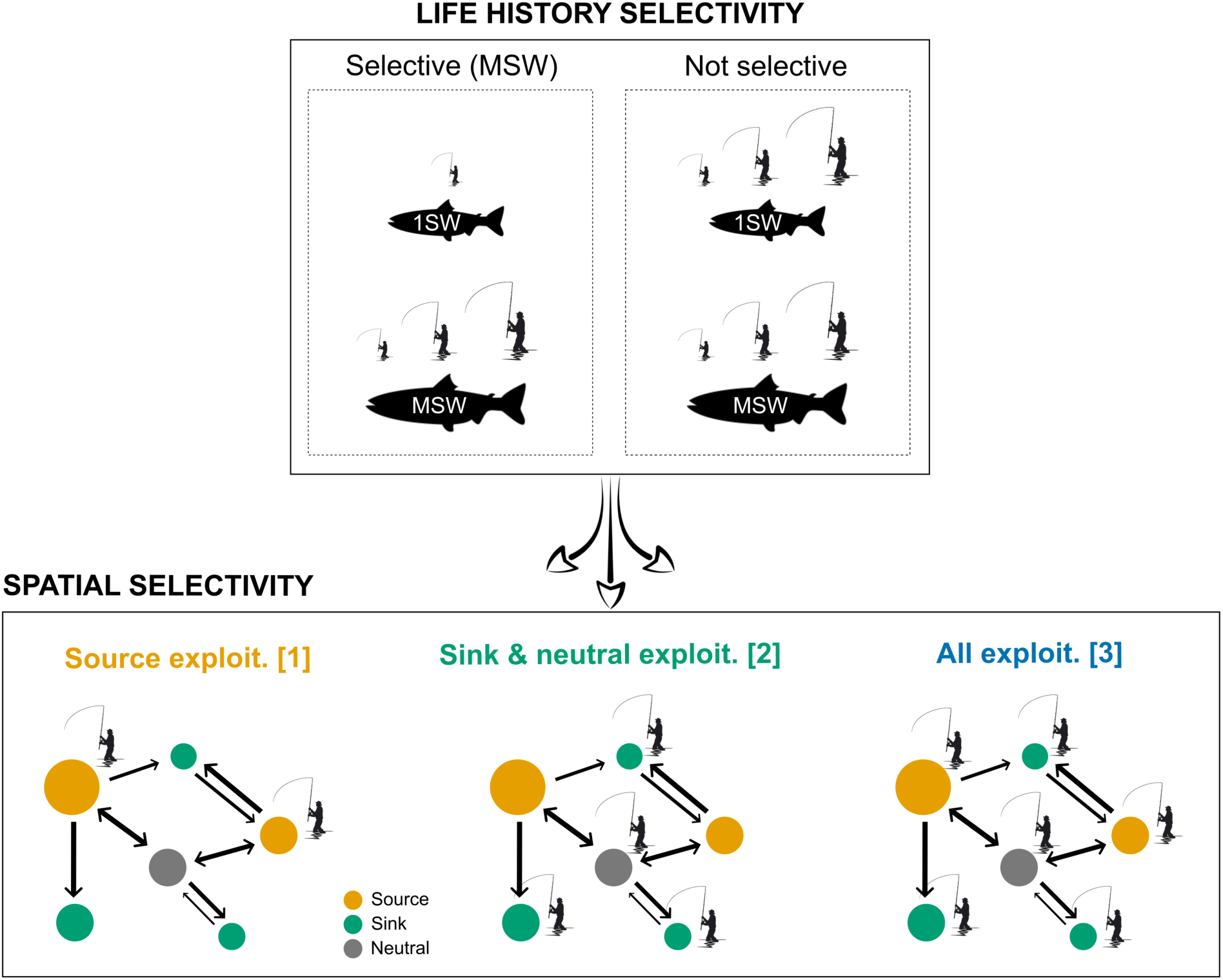
Schematic representation of the life history selectivity and spatial selectivity scenarios of exploitation, tested for a gradient of local exploitation rate from 0 to 100% and dispersal rates of 5%, 15% and 30%.

For spatially selective exploitation, we considered the interaction of the gradient of local exploitation rates, for both selective and non-selective exploitation on traits, with three management strategies (Fig. 2): two scenarios of uneven exploitation rates among populations (1 and 2) and one scenario with equal exploitation rates across the network (3). Scenarios of uneven exploitation rates among populations were based on population types, one focusing exploitation on source populations (i.e., protection of sink and neutral populations) and the other focusing on sink and neutral populations (i.e., protection of source populations). By doing so, we ensured that for each of these scenarios the exploitation was applied to approximately 50% of the total metapopulation size. Population types (sink, neutral or source, Fig. 1) were defined based on the net migration or net flow, i.e., the ratio of Immigrants/Emigrants, following the general definition of Pulliam (1988). Several underlying mechanisms can lead to variation of net migration patterns among populations and the source/sink terminology have been employed in different ways in the literature (e.g., Crowder et al., 2000; Fullerton et al., 2016; see Loreau et al., 2013 for review). In this work, asymmetries in net migration were only influenced by populations size (i.e., river catchment surface area) and the spatial configuration of the network (i.e., distance between populations). There was no difference in the intrinsic productivity among populations, and the populations we defined as “sinks” are self-sustaining in the absence of immigration at reasonable levels of exploitation.

Each combination of exploitation rate × life history selectivity × spatial selectivity was tested with dispersal rates of 5%, 15% and 30% covering the range of dispersal rates estimated in wild populations of Atlantic salmon (Jonsson et al., 2003; Consuegra et al., 2005). However for the sake of conciseness, we mainly present results for a dispersal rate of 15% as it was identified by Lamarins et al. (2022*b*) as the optimal dispersal rate for metapopulation stability in this configuration

All cross-combination scenarios (n=198) were simulated with 50 replicates each. Simulations were initialized for each population by a random draw of individuals (sampled in the same genetic and phenotypic distributions, see Lamarins et al., 2022*b*). To limit computational time, the initial number of fish drawn correspond to 25% of the river’s carrying capacity. All other parameters were kept equal for all populations (e.g., environmental conditions, trait distributions at initialization, etc.) for all scenarios. After a 10-year burn-in period with dispersal and without exploitation to allow populations stabilization, the simulations were carried out by applying treatments (i.e., exploitation rates and dispersal rates) over 40 years. This duration was sufficient to detect changes in the population dynamics and evolution of life-history traits (generation time of Atlantic salmon in the Brittany metapopulation is *∼*2.5-3.5 years, Caudal and Prévost, 2018). All simulations and analyses were performed using R version 3.6.3 and the package *metaIbasam* version 0.0.6 (https://github.com/Ibasam/MetaIBASAM). MetaIbasam code and R scripts are available at https://github.com/Ibasam/FishStrat.

### 2.3 Simulations outcomes analysis

Given the dual objective of maintaining exploitation yield and achieving conservation, we evaluated the consequences of the life history selectivity and spatial selectivity scenarios on abundance, stability, and yield at the metapopulation scale. Regarding exploitation, we averaged the total catch (i.e., summed over all populations) over the last 5 years for MSW and 1SW fish both separately and combined. Regarding metapopulation performance, we evaluated (i) the Pre-Fishery Abundance, calculated as the sum of returning adults over populations, averaged over the last 5 years; (ii) the metapopulation stability quantified as the Portfolio effect, based on the variance in abundance over the 50 years of simulation (Anderson et al., 2013; Lamarins et al., 2022*b*); and (iii) the quasi-extinction risk, computed for each population as the proportion of simulations where juvenile density dropped below a risk threshold (see Suppl. Mat.). Finally, we analysed the associated eco-evolutionary dynamics by focusing on the average traits of philopatric individuals only (i.e., excluding immigrants) over the whole metapopulation. Phenotypic changes in life history were assessed by means of the MSW/1SW ratio averaged over the last 5 years. For the underlying genetic changes, we used the male parr and female anadromous maturation thresholds and river growth potential (again averaged over the last 5 years). We computed each of these demographic, phenotypic, and genotypic metrics by averaging over simulation replicates.

To compare scenarios according to their overall exploitation intensity, the metrics described above were represented as functions of the exploitation rate measured at the scale of the metapopulation (hereafter metapopulation exploitation rate, or *MER*). This was computed *a posteriori* for each scenario by a weighted average over the last 5 years and over all simulations (Equation 1):

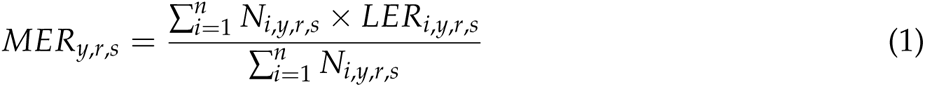

Where *y* is the year, *r* the simulation replicate, *s* the scenario, *n* the number of populations, *N_i_* the number of returns before exploitation for the population *i*, and *LER_i_* the local exploitation rate defined according to the scenario for the population *i*.

## 3 Results

We first compared the metapopulation performance, yield, and evolutionary consequences of life-history selective vs. non-selective exploitation, without any spatial selectivity. We then assessed the additional effects of the spatial variation in exploitation. It is important to note that the range of metapopulation exploitation rates (MER) depends on the type of scenarios explored. In particular, the selectivity of exploitation, whether spatially or life-history based, reduces the range of MER because captures are supported only by a fraction of the whole metapopulation (Fig. 3, 4 and 5). Moreover, when we compare the spatial scenarios for the same MER, this means that the exploitation effort on unexploited populations is transferred to the other exploited populations. Results are presented for a 15% dispersal rate; results are similar for other assumed dispersal rates (Suppl. Mat.) except as explicitly noted below.

**Figure 3:**
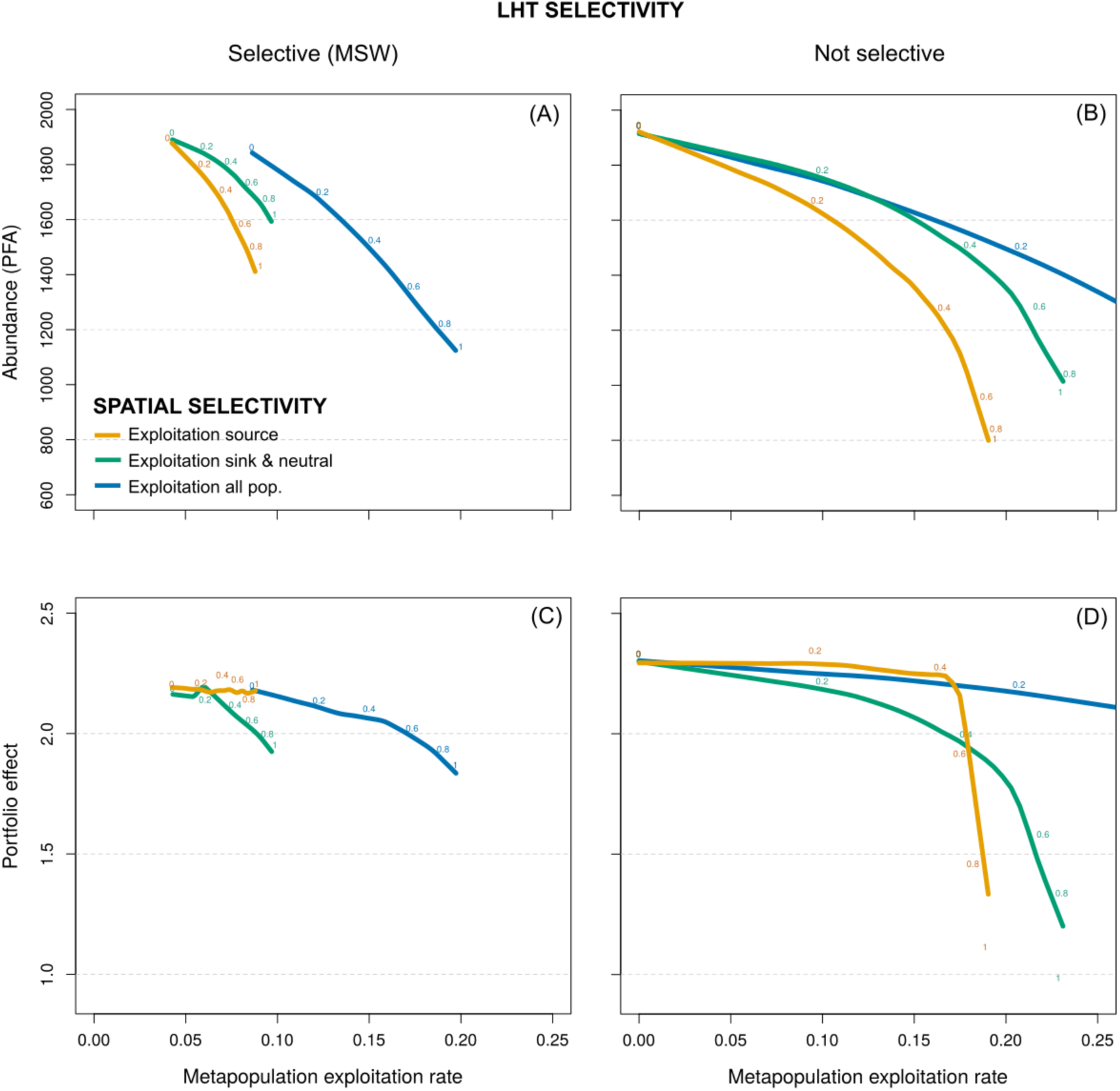
Metapopulation performance metrics averaged over the metapopulation and simulations for life history selectivity (left/right) and spatial selectivity (colors) scenarios of exploitation, for increasing metapopulation exploitation rate (MER) and a dispersal rate of 15% (other rates in Suppl. Mat.). The local exploitation rates are reported adjacent to the curves (loess regression) for each scenario. A-B) Abundance of returns (Pre-Fishery Abundance, last 5 years), C-D) Portfolio effect over the 50 years.

**Figure 4:**
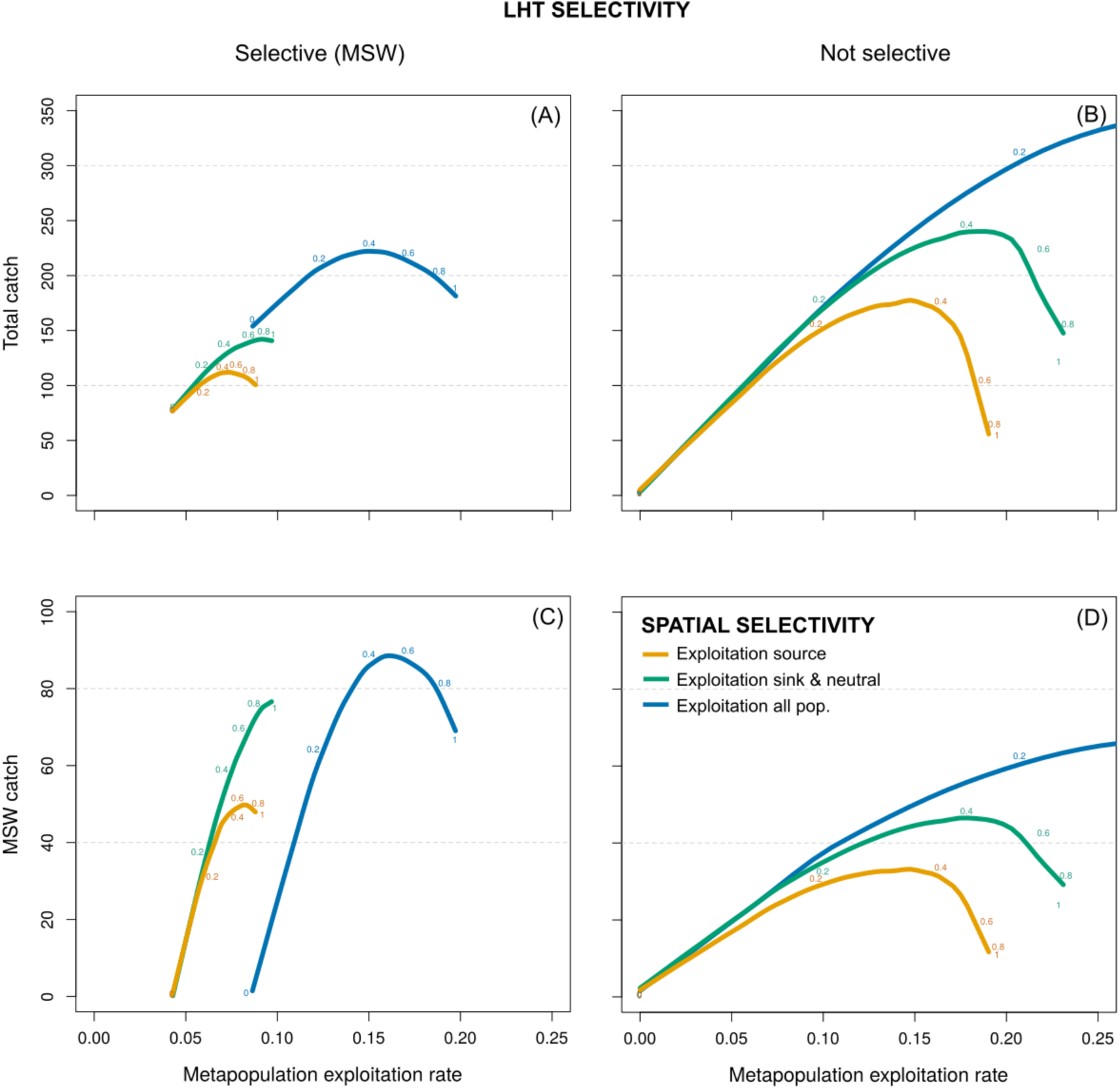
Exploitation metrics averaged over the metapopulation and simulations for life history selectivity (left/right) and spatial selectivity (colors) scenarios of exploitation, for increasing metapopulation exploitation rate (MER) and a dispersal rate of 15% (other rates in Suppl. Mat.). The local exploitation rates are reported adjacent to the curves (loess regression) for each scenario. A-B) Total catch, C-D) MSW catch (last 5 years).

**Figure 5:**
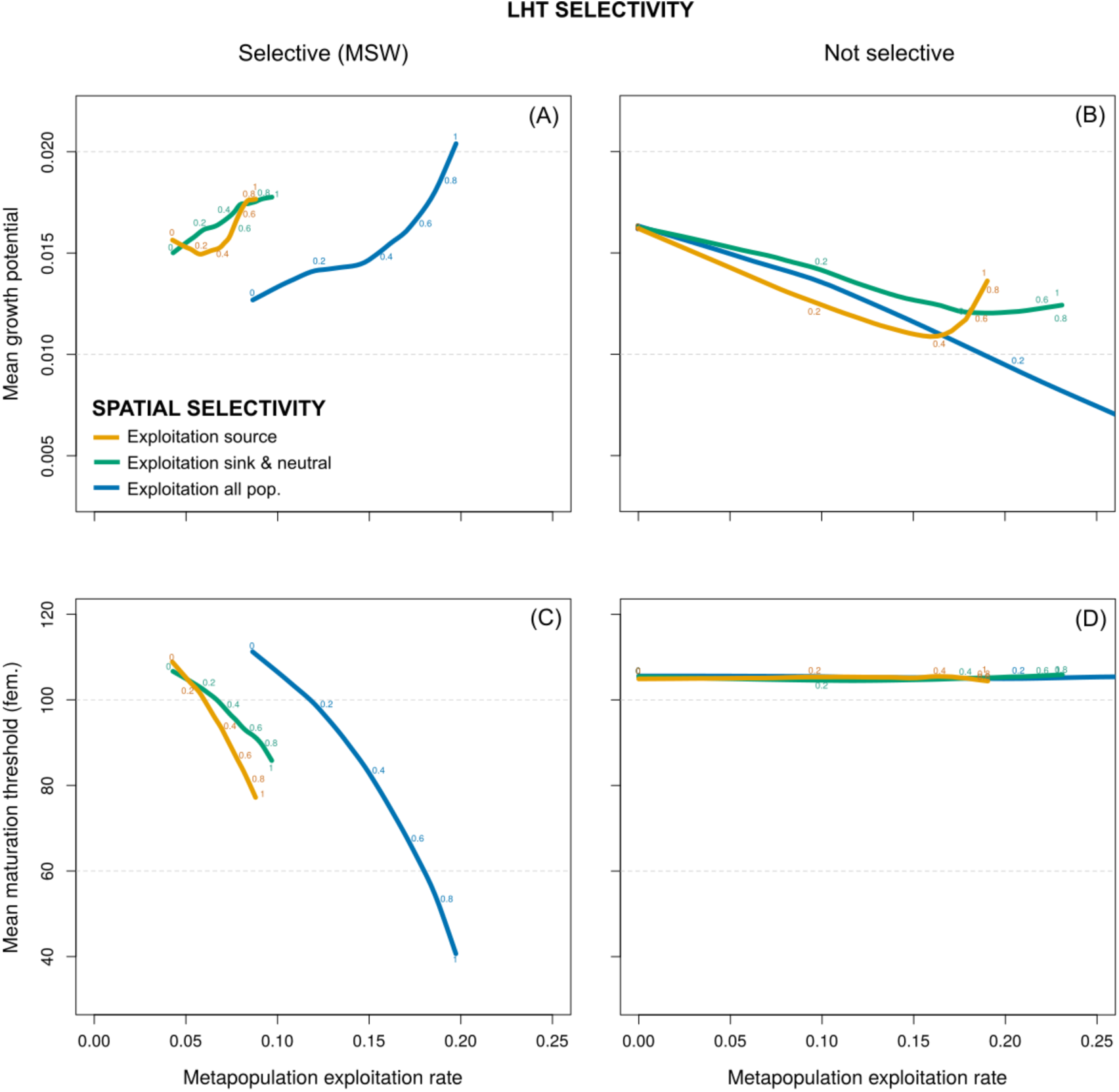
Adaptive traits (genetic) averaged over the metapopulation and simulations for life history selectivity (left/right) and spatial selectivity (colors) scenarios of exploitation, for increasing metapopulation exploitation rate (MER) and a dispersal rate of 15% (other rates in Suppl. Mat.). The local exploitation rates are reported adjacent to the curves (loess regression) for each scenario. A-B) Average genetic values of growth potential and C-D) female maturation threshold for philopatric individuals (last 5 years).

### 3.1 Life history selectivity of exploitation without spatial selectivity

#### 3.1.1 Metapopulation performance

In general, increasing the exploitation rates had a detrimental effect on metapopulation performance: it decreased both the Pre-Fishery Abundance (Fig. 3 A-B, blue) and the portfolio effect (Fig. 3 C-D, blue) showing a nonlinear accelerating decline for each, irrespective of the life-history selectivity scenario. However, the loss of metapopulation performance metrics with increasing MER was stronger in the case of selective exploitation on MSW compared to non-selective exploitation (Fig. 3 A vs B and C vs D, blue). For example, under a dispersal rate of 15%, and MER doubling from 10% to 20%, the Pre-Fishery Abundance decreased by 14% for non-selective exploitation (B), vs 37% for selective exploitation of MSW (A). Likewise, the portfolio effect decreased by 4% vs 14%. Local extinction risk of populations also increased with exploitation, and more strongly when exploitation was selective on MSW (by 100% vs 445% between 10% and 20% MER, Suppl. Mat. Fig. S4).

#### 3.1.2 Exploitation

Without selectivity, the number of MSW catches, 1SW catches, and total catches exhibited a consistent rise over the range of metapopulation exploitation rates tested (Fig. 4 B-D, blue, and Suppl. Mat. Fig. S9). But with selective exploitation of MSW, after an initial increase the MSW and the total catch peaked and then fell above a MER around 15% (Fig. 4 A-C, blue), while the number of 1SW catch continuously decreased with exploitation (Suppl. Mat. Fig. S9). Despite a higher MSW catch with selective exploitation, the total catch was lower at high MER.

#### 3.1.3 Evolution

Genetic growth potential decreased with increasing metapopulation exploitation rate in the absence of selectivity (by 32% from 10% to 20% MER with 15% of dispersal, Fig. 5 B, blue). In contrast, it increased when exploitation was selective on MSW (by 48% from 10% to 20% MER, Fig. 5 A, blue). Without selectivity, the sea maturation threshold and the MSW/1SW ratio (Fig. 5 D; Suppl. Mat. Fig. S13) remained unchanged with increasing MER, while both strongly decreased when exploitation selected against MSW fish (by 63% from 10% to 20% MER, Fig. 5 C, Suppl. Mat. Fig. S13). Even though some evolutionary changes may occur when exploitation is not selective, selection against MSW fish induced much stronger evolutionary effects, i.e., faster growth potential and lower maturation threshold, in favor of a shorter residency at sea (1SW).

### 3.2 Spatial selectivity of exploitation with or without life history selectivity

#### 3.2.1 Metapopulation performance

Regardless of life-history selectivity, the even exploitation rate across all populations resulted in the highest levels of Pre-Fishery Abundance and portfolio effect (Fig. 3, blue vs other colors, for 15% dispersal), as well as the lowest levels of average risk of local population extinction (Suppl. Mat. Fig. S4). But the relative performances of the two spatially selective exploitation strategies differed according to the dispersal rate. When the latter was high (15% or 30%), the exploitation of sink and neutral populations (protection of source populations) produced similar or lower averaged local extinction risks and higher Pre-Fishery Abundance (by 43% for MER of 20% and dispersal rate of 15%) than the exploitation of source populations, and this stands for both non-selective and selective exploitation on MSW (Fig. 3 A-B, orange vs green, Suppl. Mat. Fig. S2-S4). For lower rates of dispersal (5%), the exploitation of sink and neutral populations was no longer preferable: it resulted in levels of abundance similar to those obtained by the exploitation of source populations and it induced the highest local extinction risk (Suppl. Mat. Fig. S4). In contrast to abundance, exploiting only source populations strengthened the portfolio effect relative to focusing exploitation on sink and neutral populations (Fig. 3 C-D, orange vs green). This advantage disappeared however for high exploitation rates (*>*20%) and for a dispersal rate of 30% in the case of non-selective exploitation on life histories (Suppl. Mat. Fig. S3).

#### 3.2.2 Exploitation

Similarly, the strategy of even exploitation rate across all populations produced the highest 1SW catch and total catch with or without life-history selectivity (Fig. 4 A-B, blue vs other colors, for 15% dispersal, Suppl. Mat. Fig. S9). With non-selective exploitation on life-history, the exploitation of the source populations had the worst performance in terms of MSW, 1SW or total catch (by 77% for a 20% MER; Fig. 4 B-D, orange). The exploitation of sink and neutral populations produced intermediate results, close to those of an even exploitation rate across all populations when the dispersal rate was high (30%), and similar to those of the source exploitation with weak dispersal (5%; Suppl. Mat. Fig. S7). With life history selective exploitation, the relative performance of spatially selective strategies remained unchanged in terms of 1SW catch and total catch (Fig. 4 A, Suppl. Mat. Fig. S9). But the same MSW catch could be obtained with a lower MER with spatially selective exploitation. Thus, with a 9% MER, MSW catch 47% and 74% higher were generated under the source exploitation and the sink and neutral exploitation respectively when compared to the strategy of even exploitation rate across all populations (with a dispersal rate of 15%; Fig. 4 C).

#### 3.2.3 Evolution

Without selective exploitation of life-histories, the spatial selectivity had little evolutionary effect on the average growth potential, maturation thresholds and MSW/1SW ratio when compared to the strategy of even exploitation rate across all populations (Fig. 5 B-D, Suppl. Mat. Fig. S13). In contrast, with selective exploitation of MSW, spatial selectivity fostered an overall evolutionary response of the metapopulation. Relative to even exploitation, with spatial selectivity the growth potential evolved towards higher values by 44% and 41% (for a 9% MER and a dispersal rate of 15%) under the source and sink and neutral exploitation strategies respectively (Fig. 5 A). Similarly, the anadromous maturation threshold and the MSW/1SW ratio evolved towards lower values by 30% and 17% (for a 9% MER and a dispersal rate of 15%) under the source and sink and neutral exploitation strategies respectively. The source exploitation strategy had the strongest evolutionary effect for high dispersal rates (Fig. 5 C, Suppl. Mat. Fig. S11-S13).

## 4 Discussion

While the eco-evolutionary consequences of exploitation and dispersal are increasingly recognized (Palkovacs, 2011; Clobert et al., 2012), their joint consideration is still limited. Using a simulator of an Atlantic salmon metapopulation, we compared alternative exploitation strategies incorporating selection on life histories and on populations according to their source/sink status. Our results extended to the metapopulation scale the demographic and evolutionary effects of exploitation already found for single populations, i.e., lower abundance and earlier sexual maturation, which were stronger when exploitation was selective on larger individuals (Piou et al., 2015). The spatial selectivity of exploitation within the metapopulation based on the source/sink status of populations (based on asymmetry in net migration) had an additional detrimental effect on metapopulation performance and yield, and induced stronger evolutionary changes than when exploitation was evenly spread over all populations (Fig. 6). Non-selective exploitation, either on traits or in space, appears as the safest strategy with the dispersal rates and spatial configuration simulated. Even if obtained in a context where there was no difference in productivity among populations (population size only), this result is of special importance for management because the source/sink status of populations is essentially unknown in practice.

**Figure 6:**
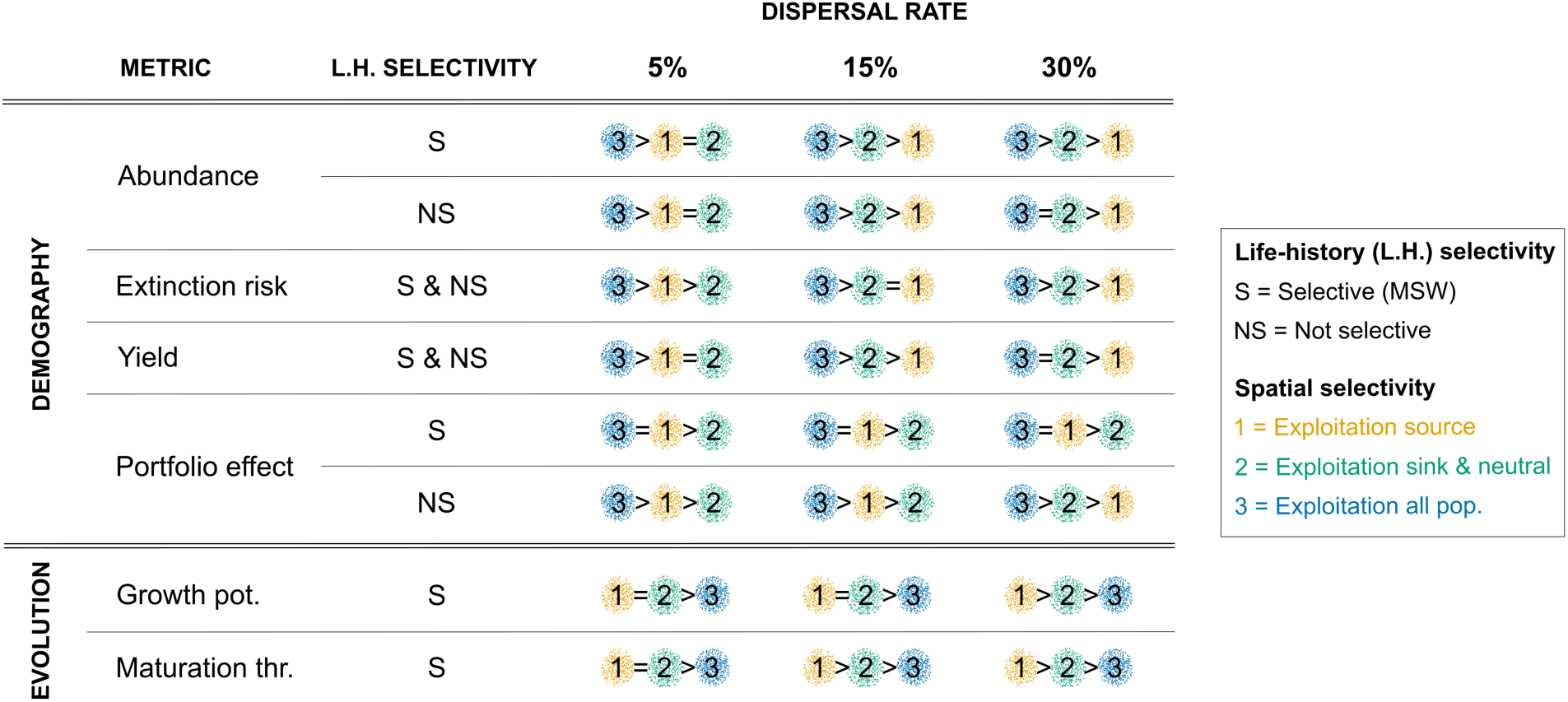
Illustrative synthesis of the relative demographic and evolutionary consequences of the three scenarios of spatial selectivity of exploitation, under both scenarios of life history selectivity and dispersal rates of 5%, 15% and 30%. Scenarios are ordered by the absolute values of demographic metrics and by the strength of evolution of the two traits.

### 4.1 Exploitation selectivity on life history

In our study, exploitation reduced the abundance and stability of the metapopulation. Using long term data, prior studies identified exploitation as a key driver of the decline of abundance and increased synchrony in population portfolios (e.g., Frank et al., 2016 in cod; Stier et al., 2020 in herring). From both empirical and simulation studies, exploitation of adults is also known to induce a faster pace of life, i.e., earlier sexual maturation at the population level (Edeline et al., 2007; Wang and Höök, 2009; Matsumura et al., 2011; Ayllón et al., 2018). In our case, lower precocious male maturation thresholds were selected for at higher exploitation rates (Suppl. Mat. Fig. S12), favoring early maturation of males in the river before seaward migration. Anadromy was thus selected against at higher exploitation levels, allowing fish to avoid the risk of being harvested before reproduction. Thus, the observed reduction in abundance and catch of anadromous adults is due to both the reduction of spawning biomass by fishing, and the evolutionary response of populations to exploitation.

Relative to non-selectivity, selective exploitation on MSW was detrimental to both metapopulation performance and yield. It further reduced the abundance and the stability of the metapopulation, increased the extinction risk of populations, and limited – or even decreased – the total catch at higher exploitation rates. Catching MSW fish disproportionately removes the fraction of the populations with a larger proportion of females with a higher fecundity and reproductive success in relation to their bigger size (Barneche et al., 2018). Hence, selective exploitation hampers the population renewal process more heavily as the number of eggs spawned by females is the initial limiting factor of the size of the next generation, the number of males being much less constraining in Atlantic salmon. Exploitation selectivity on MSW also induced the evolution of life history traits towards early breeding migration (i.e., the 1SW tactic), as a response to escape exploitation. Piou et al. (2015) obtained similar results by using a Demo-Genetic Agent-Based Model simulation of a single closed population. Our study extend these findings on the demographic and evolutionary effects of selective exploitation (Fenberg and Roy, 2008; Heino et al., 2015) to the context of a metapopulation.

### 4.2 Spatial selectivity of exploitation

Few studies have considered the consequences of spatially selective exploitation on metapopulation performance, yield, and evolution of traits in a single framework. In our work, the strategy of spatially even exploitation showed the highest levels of metapopulation performance (highest abundance, portfolio effect strength, lowest extinction risk) and yield metrics (Fig. 6), regardless of life history selectivity and the dispersal rate. Additional to demographic effects, we showed spatial selectivity of exploitation resulted in stronger evolutionary effects when fishing selected against the MSW tactic. The high local exploitation rates applied on a few populations induced evolution towards higher growth potential and lower maturation thresholds, and this was marginally mitigated by increased dispersal. Interestingly, while the even exploitation of all populations reduced the overall genetic diversity of the maturation threshold, the spatial selectivity of exploitation did the opposite. Variations in selective exploitation caused different responses among populations, inducing local adaptation and increasing overall genetic variation, especially at low dispersal rates (Suppl. Mat. Fig. S15).

We showed the relative performance of the two scenarios of selective exploitation in space may change according to dispersal intensity (Fig. 6). Indeed, compared to exploitation of source populations, the exploitation of sink and neutral populations maintained higher overall abundance, stability, reduced risk of extinction, and dampened the evolutionary effects of life history selective exploitation when dispersal was strong. This result was due to a dispersal buffering, which compensated for exploitation effects by immigration from the protected source populations to the exploited sink and neutral populations. In contrast, when exploitation targeted the source populations, their high local exploitation rates were not compensated by immigration from the sink and neutral populations. The latter would in turn be impacted by the source populations because abundance reduction and evolutionary response to exploitation would be propagated to the sink and neutral populations through immigration. However, when dispersal was weak, the exploitation of source populations appears as a better strategy as exploiting the sink and neutral populations reduced the overall abundance, stability, and increased the risk of extinction especially for small and/or isolated populations (Suppl. Mat. Fig. S5). This buffering effect of dispersal flows may relate to the “spillover” effect described in the context of Marine Protected Areas (MPAs, Di Lorenzo et al., 2016, 2020), i.e. the export of individuals outside MPAs to the surrounding fishing areas. A large body of work has shown that establishing marine reserves provides benefits for both conservation and exploitation compared to reducing fishing pressure in all habitats (e.g., Crowder et al., 2000; Rassweiler et al., 2012; Sève et al., 2023). Our results show no such benefits.

However, several factors can influence the relative performance of spatial management strategies, such as fishing effort, dispersal intensity and criteria for setting reserves (e.g. proportion, size, connectivity). Indeed, the spillover effect may not be sufficient in some cases to counteract the overexploitation of stocks outside the reserves due to displaced fishing effort (Hilborn, 2018; Ovando et al., 2021). Moreover, marine dispersal capacity (especially larval dispersal) is likely to be higher than in freshwater habitats fostering the spillover effect of MPAs. In this sense, our results showed a benefit in protecting source populations compared to the exploitation of all populations when increasing dispersal rates and for low exploitation rates. Similar to our findings, Crowder et al. (2000) showed the worst strategy was to protect the sink habitats when total fishing effort was 15%. However, they recommended protecting the source habitats for optimal population size, which contradicts our suggestion of exploiting all populations. This discrepancy might be due to different criteria for setting reserves. In their study, 20% of the habitats were designated as reserves, while our protection covered 50% of the total abundance. Additionally, the source / sink status was based on habitat quality and growth rate and dispersal was a density-dependent process in Crowder et al. (2000). Instead, we considered density-independent dispersal and equal productivity of all populations in our work. Populations intrinsic productivity has also been the criteria of populations management in other studies (Engen et al., 2018; Tufto and Hindar, 2003; Okamoto et al., 2020) that suggest to protect the smallest (mostly the sink in our case) populations to i) reduce spatial synchrony (i.e., increase the portfolio effect), ii) maximise both the total effective size and yield, and iii) reduce local populations risk of collapse, which is contrary to our results and those of Crowder et al. (2000). Altogether, these findings emphasise the difficulty of effectively managing exploited populations when the underlying processes driving their dynamics, such as dispersal, are poorly understood. Our work goes a step further by demonstrating the importance of considering the additional impacts of spatially selective exploitation on adaptive traits.

### 4.3 Lessons for management of anadromous salmonids

Our work provides novel insight into the management of anadromous salmonids (with special reference to Atlantic salmon). Our approach aimed to simultaneously consider metapopulation functioning and evolutionary perspectives while investigating the metapopulation performance, harvest and evolutionary consequences of selective exploitation based on life history traits and population types. Overall, our results advocate for an even exploitation regardless of the population or the fish trait as it provided the highest abundance, stability, yield, and the lowest extinction risk and evolutionary changes at the metapopulation scale, regardless of dispersal intensity (Fig. 6). Our results also emphasize the need to minimize selective exploitation of MSW individuals in the Atlantic salmon metapopulation, echoing the warning of MSW individuals in the Atlantic salmon metapopulation, echoing the warning of Piou et al. (2015). We additionally show that spatial selectivity of exploitation is no remedy for such selection as it tends to worsen its undesirable consequences. However, spatial selectivity induced local adaptation and fostered higher genetic variation within the population network. This is particularly interesting considering that in Atlantic salmon, (i) exploitation rate is still selective on MSW individuals and very likely varies among populations, and (ii) the source/sink status of populations is essentially unknown. Yet, it is important to point out that we primarily assessed the overall consequences of management strategies at the metapopulation scale. Our results are more contrasted at the local population scale depending on dispersal intensity and spatial configuration. The even exploitation of the metapopulation and conservation of source populations can still put at risk local populations, especially if they are small and isolated (Suppl. Mat. Fig. S5). Whether this local variation in performance is significant or not should also be considered by managers.

Because it offers opportunities for influencing rescue effects or just of constraining the upper range of metapopulation exploitation rate, spatial selectivity of exploitation based on the source/sink status of the populations might have appeared intuitively appealing. The protection of part of the metapopulation could be an easy way to implement and control the maximum rate of exploitation. Setting fishing reserves for some populations and letting fisheries occur (even with little constraints) on the remainder might be more straightforward than regulating the local exploitation rate of each population. According to our study, this might be relevant, both in terms of conservation and yield, provided that (i) reserves are set for the source populations, (ii) the metapopulation exploitation rate is not too high, (iii) exploitation is not selective on life histories, and (iv) the dispersal rate is high. These conditions are not easily met in practice because fishers have a preference for catching the large MSW fish, the source/sink status of populations is essentially unknown and limiting the metapopulation exploitation rate may require regulation of the local exploitation rates on the sink populations. The latter point might be circumvented by concentrating exploitation on a small fraction of the metapopulation (i.e., a small number of sink systems) but this may face acceptance issues among the fishing community. In addition, beyond the uncertainty of the source/sink status of populations, the status can potentially change over time due to the internal dynamics of the metapopulation or to external factors, such as exploitation itself, as discussed by Lin et al. (2011). Hence, there is a risk associated with the implementation of a spatially selective strategy, which cannot readily be controlled and remains to be quantified.

Our recommendation for an even exploitation of a metapopulation over all its spatial or phenotypic components remains contingent on some important founding assumptions. First, we modeled dispersal as density independent and based on an isolation by distance mechanism. Although this appears reasonable for a long-range migratory species with a philopatric behaviour like Atlantic salmon, it remains hypothetical (e.g., Pearse et al., 2011) as the proximate causes of dispersal remain unknown. Second, there is no variation in productivity among populations in our setting. This seems sensible when the metapopulation is distributed over an area quite homogeneous environmentally, like the western part of the Brittany peninsula which holds the Atlantic salmon metapopulation we simulated. If not verified, the literature on mixed stock fisheries (Ricker, 1958; Ying et al., 2011) shows that the least productive populations can be threatened by an even exploitation of a metapopulation. Finally, the specific spatial configuration we simulated, inspired from the Brittany metapopulation, may also influence our results. The relative proportion of source and sink populations (and thus the proportion of total abundance under protection), along with their spatial distribution across the network, may drive the demographic and evolutionary responses of the network to exploitation. These factors have been shown to affect the adaptive capacity and demographic recovery of populations following a perturbation (Lamarins et al., 2024). We therefore advise managers to critically assess the relevance of our assumptions in the specific cases they may have to deal with.

### 4.4 Perspectives

Demo-Genetic Agent-Based Models are useful for exploring eco-evolutionary consequences of alternative management strategies that are challenging to test in nature. In our case, MetaIBASAM allowed the emergence of varying demographic and evolutionary outcomes depending on which life history was selected or which type of population was exploited. But our results are conditional on some important assumptions that are debatable and could be changed or relaxed by further work. First, for the sake of parsimony, we assumed dispersal (i.e., emigration) rate was constant over space and time and across individuals. However, empirical studies reported variability over space (Consuegra et al., 2005) and time (Jonsson et al., 2003) in Atlantic salmon, and suggested that dispersal propensity might depend on individual traits (e.g., genotype, Saastamoinen et al., 2018) or population features (e.g., density-dependence, Berdahl et al., 2016). Taking into account these results in our modeling might result in different outcomes. For example, with a genetic basis, one might expect dispersal rates to evolve as function of the spatial dimension of exploitation, and exploitation targeted on sink populations might select for higher dispersal in these populations but lower in source populations, and vice-versa.

We also evaluated the consequences of management strategies that were contrasted by setting protected areas (no harvest at all) on source versus sink populations, with very high local exploitation rates in some instances. We thus defined our strategies based on a single dimension of populations diversity: whether they are source populations or demographically sink, which was only dependent on population size and spatial arrangement (proximity to nearby populations). This work should be extended to investigate different spatial configurations, various proportions of total abundance or habitats under conservation, and varying dispersal intensities. Differences in other dimensions of biodiversity such as effective population size (Hindar et al., 2004), evenness, asynchrony in dynamics, heterogeneity in stock productivity (Moore et al., 2021), environmental, or genetic features should also be considered in future work. Many of these population-specific features are and will most likely remain essentially unknown to managers though. The challenge for managers is to choose strategies which are robust to the uncertainties they face. Simulations using DG-ABMs offer very interesting perspectives for assessing this robustness by means of experimental designs varying both local population features and spatial allocation of fishing rules.

Finally, we simulated a context of stationarity of the environment, i.e., no change over time of the mean environmental conditions. Yet, climate change will be an additional selective pressure. Its interaction with exploitation has received some attention (Piou et al., 2015), but not in a metapopulation context. As discussed by Bradford and Braun (2021), protecting theproductive source populations might help to maintain the abundance of a metapopulation in the short term. In the longer term, preserving a diversity of habitats and diversified populations (phenotypically and genetically), could prove relevant. This biocomplexity fosters resilience in metapopulations confronting environmental changes (Anderson et al., 2015; Walsworth et al., 2019; Moore and Schindler, 2022), and stability of yields in exploited species (Hilborn et al., 2003; Schindler et al., 2010; Freshwater et al., 2019; Connors et al., 2022). Managing a network of diversified populations may prove crucial to foster the stability and the resilience of a metapopulation facing unpredictable future, and our work emphasizes the need to consider dispersal as part of the overall equation influencing population diversity.

## Acknowledgments

We thank Tewann Beauchard for helping with preliminary scenarios analysis. We also want to thank the Bretagne Grands Migrateurs association and angling clubs for allowing us access to Atlantic salmon datasets from Brittany. We gratefully acknowledge funding from the Region Nouvelle Aquitaine, E2S-UPPA, INRAE and the French Biodiversity Agency (OFB) via the unit Management of Diadromous Fish in their Environment (pôle MIAME). This work was conducted within the International Associated Laboratory MacLife.

## Data availability

Modeling code and R scripts are freely available at https://github.com/Ibasam/FishStrat.

## Literature Cited

1. Aas, Ø., A. Klemetsen, S. Einum, and J. Skurdal. 2010. Atlantic Salmon Ecology. John Wiley & Sons.

2. Akçakaya, H. R., G. Mills, and C. P. Doncaster. 2007. The role of metapopulations in conservation. Pages 64–84 in D. W. Macdonald and K. Service, eds. Key Topics in Conservation Biology. Blackwell Publishing.

3. Allendorf, F. W., and J. J. Hard. 2009. Human-induced evolution caused by unnatural selection through harvest of wild animals. Proceedings of the National Academy of Sciences 106:9987– 9994.

4. Anderson, S. C., A. B. Cooper, and N. K. Dulvy. 2013. Ecological prophets: Quantifying metapopulation portfolio effects. Methods in Ecology and Evolution 4:971–981.

5. Anderson, S. C., J. W. Moore, M. M. McClure, N. K. Dulvy, and A. B. Cooper. 2015. Portfolio conservation of metapopulations under climate change. Ecological Applications 25:559–572.

6. Ayllón, D., G. G. Nicola, B. Elvira, and A. Almodóvar. 2019. Optimal harvest regulations under conflicting tradeoffs between conservation and recreational fishery objectives. Fisheries Research 216:47–58.

7. Ayllón, D., S. F. Railsback, A. Almodóvar, G. G. Nicola, S. Vincenzi, B. Elvira, and V. Grimm. 2018. Eco-evolutionary responses to recreational fishing under different harvest regulations. Ecology and Evolution 8:9600–9613.

8. Barneche, D. R., D. R. Robertson, C. R. White, and D. J. Marshall. 2018. Fish reproductive-energy output increases disproportionately with body size. Science 360:642–645.

9. Benson, A. J., S. P. Cox, and J. S. Cleary. 2015. Evaluating the conservation risks of aggregate harvest management in a spatially-structured herring fishery. Fisheries Research 167:101–113.

10. Berdahl, A., P. A. H. Westley, S. A. Levin, I. D. Couzin, and T. P. Quinn. 2016. A collective navigation hypothesis for homeward migration in anadromous salmonids. Fish and Fisheries 17:525–542.

11. Bouchard, C., M. Buoro, C. Lebot, and S. M. Carlson. 2022. Synchrony in population dynamics of juvenile Atlantic salmon: Analyzing spatiotemporal variation and the influence of river flow and demography. Canadian Journal of Fisheries and Aquatic Sciences 79:782–794.

12. Bradbury, I. R., L. C. Hamilton, S. Rafferty, D. Meerburg, R. Poole, J. B. Dempson, M. J. Robertson, D. G. Reddin, V. Bourret, M. Dionne, G. Chaput, T. F. Sheehan, T. L. King, J. R. Candy, and L. Bernatchez. 2015. Genetic evidence of local exploitation of Atlantic salmon in a coastal subsistence fishery in the Northwest Atlantic. Canadian Journal of Fisheries and Aquatic Sciences 72:83–95.

13. Bradford, M. J., and D. C. Braun. 2021. Regional and local effects drive changes in spawning stream occupancy in a sockeye salmon metapopulation. Canadian Journal of Fisheries and Aquatic Sciences 78:1084–1095.

14. Brown, J. H., and A. Kodric-Brown. 1977. Turnover rates in insular biogeography: Effect of immigration on extinction. Ecology 58:445–449.

15. Carlson, S. M., C. J. Cunningham, and P. A. H. Westley. 2014. Evolutionary rescue in a changing world. Trends in Ecology & Evolution 29:521–530.

16. Caudal, A.-L., and E. Prévost. 2018. Bilan du suivi du stock de saumon sur le Scorff. Synthèse 1994-2017. Tech. rep., Fédération du Morbihan Pour la Pêche et la Protection du Milieu Aquatique.

17. Chaput, G. 2012. Overview of the status of Atlantic salmon (*Salmo salar*) in the North Atlantic and trends in marine mortality. ICES Journal of Marine Science 69:1538–1548.

18. Clobert, J., M. Baguette, T. G. Benton, and J. M. Bullock. 2012. Dispersal Ecology and Evolution. Oxford University Press.

19. Clobert, J., E. Danchin, A. A. Dhondt, and J. D. Nichols, eds. 2001. Dispersal. Oxford University Press, Oxford, New York.

20. Connors, B. M., M. R. Siegle, J. Harding, S. Rossi, B. A. Staton, M. L. Jones, M. J. Bradford, R. Brown, B. Bechtol, B. Doherty, S. Cox, and B. J. G. Sutherland. 2022. Chinook salmon diversity contributes to fishery stability and trade-offs with mixed-stock harvest. Ecological Applications n/a:e2709.

21. Consuegra, S., E. Verspoor, D. Knox, and C. García de Leániz. 2005. Asymmetric gene flow and the evolutionary maintenance of genetic diversity in small, peripheral Atlantic salmon populations. Conservation Genetics 6:823–842.

22. Cooper, A. B., and M. Mangel. 1999. The dangers of ignoring metapopulation structure for the conservation of salmonids. Fishery Bulletin-National Oceanic and Atmospheric Administration 97:213–226.

23. Crowder, L. B., S. J. Lyman, W. F. Figueira, and J. Priddy. 2000. Source-sink population dynamics and the problem of siting marine reserves. BULLETIN OF MARINE SCIENCE 66:22.

24. Czorlich, Y., T. Aykanat, J. Erkinaro, P. Orell, and C. R. Primmer. 2022. Rapid evolution in salmon life history induced by direct and indirect effects of fishing. Science 376:420–423.

25. Dempson, J. B., C. J. Schwarz, D. G. Reddin, M. F. O’Connell, C. C. Mullins, and C. E. Bourgeois. 2001. Estimation of marine exploitation rates on Atlantic salmon (*Salmo salar* L.) stocks in Newfoundland, Canada. ICES Journal of Marine Science 58:331–341.

26. Di Lorenzo, M., J. Claudet, and P. Guidetti. 2016. Spillover from marine protected areas to adjacent fisheries has an ecological and a fishery component. Journal for Nature Conservation 32:62–66.

27. Di Lorenzo, M., P. Guidetti, A. Di Franco, A. Calò, and J. Claudet. 2020. Assessing spillover from marine protected areas and its drivers: A meta-analytical approach. Fish and Fisheries 21:906–915.

28. Edeline, E., S. M. Carlson, L. C. Stige, I. J. Winfield, J. M. Fletcher, J. B. James, T. O. Haugen, L. A. Vøllestad, and N. C. Stenseth. 2007. Trait changes in a harvested population are driven by a dynamic tug-of-war between natural and harvest selection. Proceedings of the National Academy of Sciences 104:15799–15804.

29. Enberg, K., C. Jørgensen, E. S. Dunlop, M. Heino, and U. Dieckmann. 2009. Implications of fisheries-induced evolution for stock rebuilding and recovery. Evolutionary Applications 2:394– 414.

30. Engen, S., F. J. Cao, and B.-E. Sæther. 2018. The effect of harvesting on the spatial synchrony of population fluctuations. Theoretical Population Biology 123:28–34.

31. Erkinaro, J., *, Y. Czorlich, *, P. Orell, J. Kuusela, M. Falkegård, M. Länsman, H. Pulkkinen, C. R. Primmer, and E. Niemelä. 2019. Life history variation across four decades in a diverse population complex of Atlantic salmon in a large subarctic river. Canadian Journal of Fisheries and Aquatic Sciences 76:42–55.

32. Fenberg, P. B., and K. Roy. 2008. Ecological and evolutionary consequences of size-selective harvesting: How much do we know? Molecular Ecology 17:209–220.

33. Frank, K. T., B. Petrie, W. C. Leggett, and D. G. Boyce. 2016. Large scale, synchronous variability of marine fish populations driven by commercial exploitation. Proceedings of the National Academy of Sciences 113:8248–8253.

34. Freshwater, C., S. C. Anderson, K. R. Holt, A.-M. Huang, and C. A. Holt. 2019. Weakened portfolio effects constrain management effectiveness for population aggregates. Ecological Applications 29:e01966.

35. Fullerton, A. H., S. Anzalone, P. Moran, D. M. Van Doornik, T. Copeland, and R. W. Zabel. 2016. Setting spatial conservation priorities despite incomplete data for characterizing metapopulations. Ecological Applications 26:2560–2580.

36. Garcia, S. M., J. Kolding, J. Rice, M.-J. Rochet, S. Zhou, T. Arimoto, J. E. Beyer, L. Borges, A. Bundy, D. Dunn, E. A. Fulton, M. Hall, M. Heino, R. Law, M. Makino, A. D. Rijnsdorp, F. Simard, and A. D. M. Smith. 2012. Reconsidering the consequences of selective fisheries. Science 335:1045– 1047.

37. Govaert, L., E. A. Fronhofer, S. Lion, C. Eizaguirre, D. Bonte, M. Egas, A. P. Hendry, A. De Brito Martins, C. J. Melián, J. A. M. Raeymaekers, I. I. Ratikainen, B.-E. Saether, J. A. Schweitzer, and B. Matthews. 2019. Eco-evolutionary feedbacks—Theoretical models and perspectives. Functional Ecology 33:13–30.

38. Grimm, V., E. Revilla, U. Berger, F. Jeltsch, W. M. Mooij, S. F. Railsback, H.-H. Thulke, J. Weiner, T. Wiegand, and D. L. DeAngelis. 2005. Pattern-oriented modeling of agent-based complex systems: Lessons from ecology. Science (New York, N.Y.) 310:987–991.

39. Heino, M., B. Díaz Pauli, and U. Dieckmann. 2015. Fisheries-induced evolution. Annual Review of Ecology, Evolution, and Systematics 46:461–480.

40. Hilborn, R. 2018. Are MPAs effective? ICES Journal of Marine Science 75:1160–1162.

41. Hilborn, R., T. P. Quinn, D. E. Schindler, and D. E. Rogers. 2003. Biocomplexity and fisheries sustainability. Proceedings of the National Academy of Sciences 100:6564–6568.

42. Hindar, K., J. Tufto, L. M. Sættem, and T. Balstad. 2004. Conservation of genetic variation in harvested salmon populations. ICES Journal of Marine Science 61:1389–1397.

43. Hoffmann, A., P. Griffin, S. Dillon, R. Catullo, R. Rane, M. Byrne, R. Jordan, J. Oakeshott, A. Weeks, L. Joseph, P. Lockhart, J. Borevitz, and C. Sgrò. 2015. A framework for incorporating evolutionary genomics into biodiversity conservation and management. Climate Change Responses 2:1.

44. Hutchinson, W. F. 2008. The dangers of ignoring stock complexity in fishery management: The case of the North Sea cod. Biology Letters 4:693–695.

45. ICES. 2021. Working group on North Atlantic salmon. Scientific Report 3 (9), ICES-CIEM.

46. Jonsson, B., N. Jonsson, and L. P. Hansen. 2003. Atlantic salmon straying from the River Imsa. Journal of Fish Biology 62:641–657.

47. Kininmonth, S., R. Weeks, R. A. Abesamis, L. P. C. Bernardo, M. Beger, E. A. Treml, D. Williamson, and R. L. Pressey. 2019. Strategies in scheduling marine protected area establishment in a network system. Ecological Applications 29:e01820.

48. Kuparinen, A., and J. Merilä. 2007. Detecting and managing fisheries-induced evolution. Trends in Ecology & Evolution 22:652–659.

49. Lamarins, A., V. Fririon, D. Folio, C. Vernier, L. Daupagne, J. Labonne, M. Buoro, F. Lefèvre, C. Piou, and S. Oddou-Muratorio. 2022a. Importance of interindividual interactions in eco-evolutionary population dynamics: The rise of demo-genetic agent-based models. Evolutionary Applications 15:1988–2001.

50. Lamarins, A., F. Hugon, C. Piou, J. Papaïx, E. Prévost, S. M. Carlson, and M. Buoro. 2022b. Implications of dispersal in Atlantic salmon: Lessons from a demo-genetic agent-based model. Canadian Journal of Fisheries and Aquatic Sciences 79:2025–2042.

51. Lamarins, A., E. Prévost, S. M. Carlson, and M. Buoro. 2024. The importance of network spatial structure as a driver of eco-evolutionary dynamics. Ecography 2024:e06933.

52. Lebot, C., M.-A. Arago, L. Beaulaton, G. Germis, M. Nevoux, E. Rivot, and E. Prévost. 2022. Taking full advantage of the diverse assemblage of data at hand to produce time series of abundance: A case study on Atlantic salmon populations of Brittany. Canadian Journal of Fisheries and Aquatic Sciences 79:533–547.

53. Lin, J. E., R. Hilborn, T. P. Quinn, and L. Hauser. 2011. Self-sustaining populations, population sinks or aggregates of strays: Chum (*Oncorhynchus keta*) and Chinook salmon (*Oncorhynchus tshawytscha*) in the Wood River system, Alaska. Molecular Ecology 20:4925–4937.

54. Loreau, M., T. Daufresne, A. Gonzalez, D. Gravel, F. Guichard, S. J. Leroux, N. Loeuille, F. Massol, and N. Mouquet. 2013. Unifying sources and sinks in ecology and Earth sciences. Biological Reviews 88:365–379.

55. Marty, L., U. Dieckmann, and B. Ernande. 2015. Fisheries-induced neutral and adaptive evolution in exploited fish populations and consequences for their adaptive potential. Evolutionary Applications 8:47–63.

56. Matsumura, S., R. Arlinghaus, and U. Dieckmann. 2011. Assessing evolutionary consequences of size-selective recreational fishing on multiple life-history traits, with an application to northern pike (*Esox lucius*). Evolutionary Ecology 25:711–735.

57. Moore, J. W., B. M. Connors, and E. E. Hodgson. 2021. Conservation risks and portfolio effects in mixed-stock fisheries. Fish and Fisheries 22:1024–1040.

58. Moore, J. W., and D. E. Schindler. 2022. Getting ahead of climate change for ecological adaptation and resilience. Science 376:1421–1426.

59. Okamoto, D. K., M. Hessing-Lewis, J. F. Samhouri, A. O. Shelton, A. Stier, P. S. Levin, and A. K. Salomon. 2020. Spatial variation in exploited metapopulations obscures risk of collapse. Ecological Applications 30:e02051.

60. Olmos, M., F. Massiot-Granier, E. Prévost, G. Chaput, I. R. Bradbury, M. Nevoux, and E. Rivot. 2019. Evidence for spatial coherence in time trends of marine life history traits of Atlantic salmon in the North Atlantic. Fish and Fisheries 20:322–342.

61. Olsen, E. M., M. Heino, G. R. Lilly, M. J. Morgan, J. Brattey, B. Ernande, and U. Dieckmann. 2004. Maturation trends indicative of rapid evolution preceded the collapse of northern cod. Nature 428:932–935.

62. Ovando, D., J. E. Caselle, C. Costello, O. Deschenes, S. D. Gaines, R. Hilborn, and O. Liu. 2021. Assessing the population-level conservation effects of marine protected areas. Conservation Biology 35:1861–1870.

63. Palkovacs, E. P. 2011. The overfishing debate: An eco-evolutionary perspective. Trends in Ecology & Evolution 26:616–617.

64. Pearse, D. E., E. Martinez, and J. C. Garza. 2011. Disruption of historical patterns of isolation by distance in coastal steelhead. Conservation Genetics 12:691–700.

65. Pelletier, D., and S. Mahévas. 2005. Spatially explicit fisheries simulation models for policy evaluation. Fish and Fisheries 6:307–349.

66. Perrier, C., R. Guyomard, J.-L. Bagliniere, and G. Evanno. 2011. Determinants of hierarchical genetic structure in Atlantic salmon populations: Environmental factors vs. anthropogenic influences. Molecular Ecology 20:4231–4245.

67. Piou, C., and E. Prévost. 2012. A demo-genetic individual-based model for Atlantic salmon populations: Model structure, parameterization and sensitivity. Ecological Modelling 231:37– 52.

68. Piou, C., and E. Prévost. 2013. Contrasting effects of climate change in continental vs. oceanic environments on population persistence and microevolution of Atlantic salmon. Global Change Biology 19:711– 723.

69. Piou, C., M. H. Taylor, J. Papaix, and E. Prevost. 2015. Modelling the interactive effects of selective fishing and environmental change on Atlantic salmon demogenetics. Journal of Applied Ecology 52:1629–1637.

70. Pulliam, H. R. 1988. Sources, sinks, and population regulation. The American Naturalist 132:652– 661.

71. Radici, A., J. Claudet, A. Ligas, I. Bitetto, G. Lembo, M. T. Spedicato, P. Sartor, C. Piccardi, and P. Melià. 2023. Assessing fish–fishery dynamics from a spatially explicit metapopulation perspective reveals winners and losers in fisheries management. Journal of Applied Ecology 60:2482–2493.

72. Rassweiler, A., C. Costello, and D. A. Siegel. 2012. Marine protected areas and the value of spatially optimized fishery management. Proceedings of the National Academy of Sciences 109:11884–11889.

73. Ricker, W. E. 1958. Maximum sustained yields from fluctuating environments and mixed stocks. Journal of the Fisheries Research Board of Canada 15:991–1006.

74. Saastamoinen, M., G. Bocedi, J. Cote, D. Legrand, F. Guillaume, C. W. Wheat, E. A. Fronhofer, C. Garcia, R. Henry, A. Husby, M. Baguette, D. Bonte, A. Coulon, H. Kokko, E. Matthysen, K. Niitepõld, E. Nonaka, V. M. Stevens, J. M. J. Travis, K. Donohue, J. M. Bullock, and M. del Mar Delgado. 2018. Genetics of dispersal. Biological Reviews 93:574–599.

75. Schindler, D. E., R. Hilborn, B. Chasco, C. P. Boatright, T. P. Quinn, L. A. Rogers, and M. S. Webster. 2010. Population diversity and the portfolio effect in an exploited species. Nature 465:609–612.

76. Schtickzelle, N., and T. P. Quinn. 2007. A metapopulation perspective for salmon and other anadromous fish. Fish and Fisheries 8:297–314.

77. Sève, C., M. Belharet, P. Melià, A. Di Franco, A. Calò, and J. Claudet. 2023. Fisheries outcomes of marine protected area networks: Levels of protection, connectivity, and time matter. Conservation Letters 16:e12983.

78. Smith, P. J., R. I. C. C. Francis, and M. McVeagh. 1991. Loss of genetic diversity due to fishing pressure. Fisheries Research 10:309–316.

79. Stier, A. C., A. Olaf Shelton, J. F. Samhouri, B. E. Feist, and P. S. Levin. 2020. Fishing, environment, and the erosion of a population portfolio. Ecosphere 11:e03283.

80. Thériault, V., E. S. Dunlop, U. Dieckmann, L. Bernatchez, and J. J. Dodson. 2008. The impact of fishing-induced mortality on the evolution of alternative life-history tactics in brook charr. Evolutionary Applications 1:409–423.

81. Thorley, J. L., A. F. Youngson, and R. Laughton. 2007. Seasonal variation in rod recapture rates indicates differential exploitation of Atlantic salmon, *Salmo salar*, stock components. Fisheries Management and Ecology 14:191–198.

82. Tufto, J., and K. Hindar. 2003. Effective size in management and conservation of subdivided populations. Journal of Theoretical Biology 222:273–281.

83. Walsworth, T. E., D. E. Schindler, M. A. Colton, M. S. Webster, S. R. Palumbi, P. J. Mumby, T. E. Essington, and M. L. Pinsky. 2019. Management for network diversity speeds evolutionary adaptation to climate change. Nature Climate Change 9:632–636.

84. Wang, H.-Y., and T. O. Höök. 2009. Eco-genetic model to explore fishing-induced ecological and evolutionary effects on growth and maturation schedules. Evolutionary Applications 2:438– 455.

85. Ying, Y., Y. Chen, L. Lin, and T. Gao. 2011. Risks of ignoring fish population spatial structure in fisheries management. Canadian Journal of Fisheries and Aquatic Sciences 68:2101–2120.

86. Zurell, D., C. König, A.-K. Malchow, S. Kapitza, G. Bocedi, J. Travis, and G. Fandos. 2022. Spatially explicit models for decision-making in animal conservation and restoration. Ecography 2022.

