## Supplementary Figures for "Eco-Evolutionary Consequences of Selective Exploitation on Metapopulations Illustrated With Atlantic Salmon"

### Appendix S: Supplementary Materials

#### 4.5 Model overview

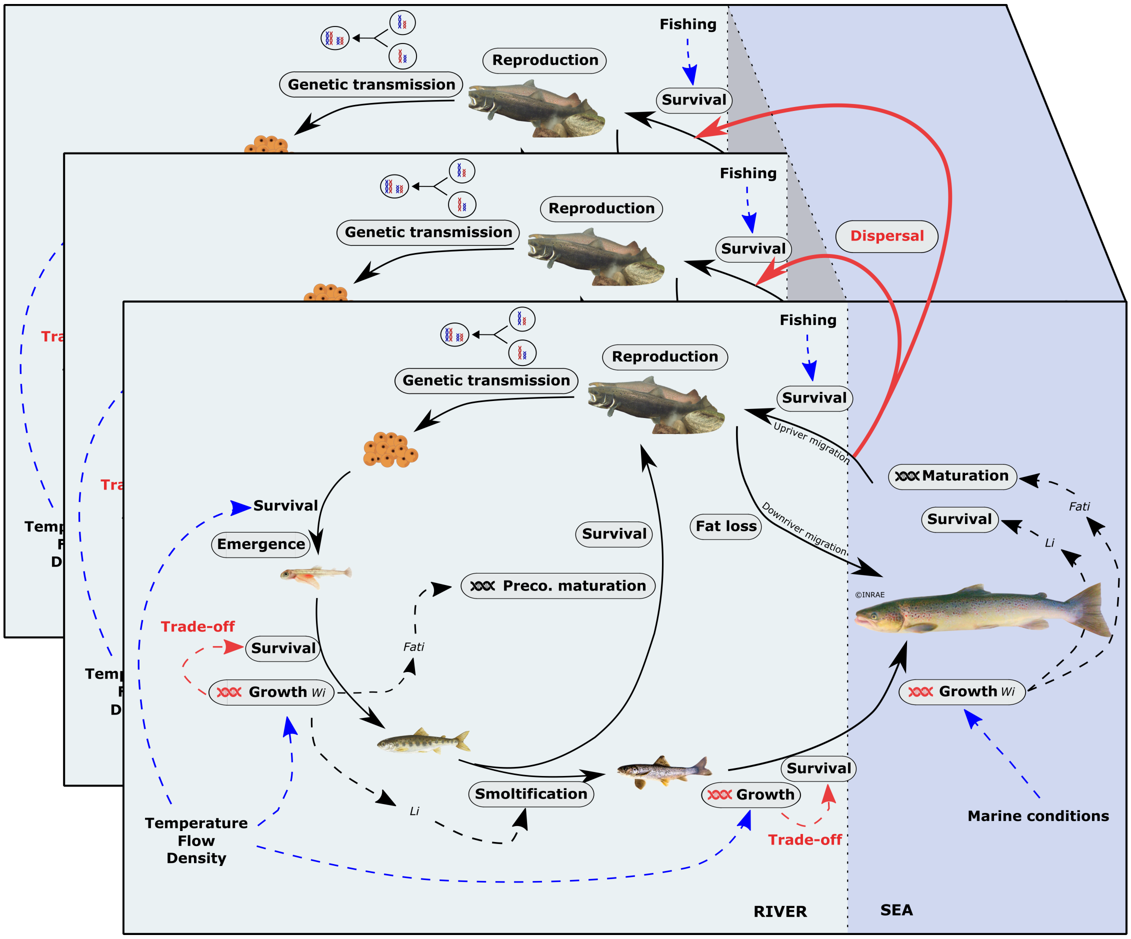


Figure S1: Schematic overview of the MetaIBASAM model, from Lamarins et al. (2022*b*). Processes at individual levels are highlighted in gray, where the DNA icon indicates heritable traits linked to these processes (maturation thresholds and growth potential). The dashed arrows represent the influence of both environmental and anthropogenic factors (in blue), or the influence of state variables of individuals (in italics). Each big rectangle represents a population, exchanging individuals with neighboring populations via the dispersal process.

In this Supplementary Materials, we present the model overview, and the results for all metrics and all scenarios, i.e., exploitation rates, selectivity based on traits, spatial selectivity, and dispersal rates.

#### 4.6 Metapopulation performance metrics: abundance

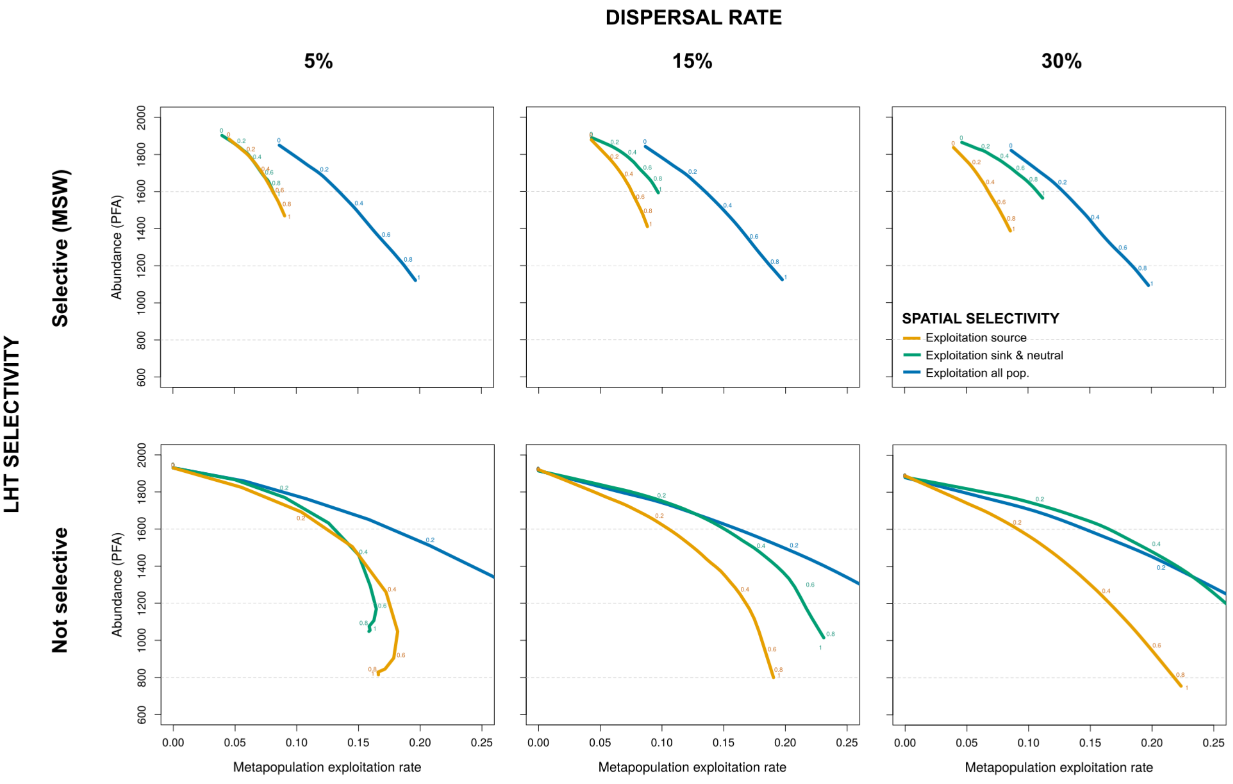


Figure S2: Metapopulation abundance (Pre-Fishery Abundance, last 5 years) averaged over simulations for each scenario of dispersal rate (5%, 15%, 30%), life history selectivity (top/bottom) and spatial selectivity (colors), for increasing metapopulation exploitation rate (MER). The local exploitation rates are reported adjacent to the curves (loess regression) for each scenario.

The Pre-Fishery Abundance (hereafter *PFA*) was calculated as the sum of returning adults over populations averaged over the last 5 years and over simulation replicates.

*Effect of life history selectivity.* Regardless of dispersal rate and spatial selectivity scenarios, the PFA was lower for increasing exploitation rates and when exploitation was selective on MSW fish.

*Effect of spatial selectivity.* Overall, the PFA was the highest for the strategy of even exploitation rate across all populations regardless of the selectivity scenarios and dispersal rate. For a low dispersal rate, the two spatial exploitation strategies showed similar levels of PFA, while the strategy of exploitation of sink and neutral populations showed higher PFA than the strategy of source exploitation for higher dispersal rates. It even reached similar levels of PFA than the strategy of even exploitation rates for a dispersal rate of 30% and non-selective exploitation on traits.

#### 4.7 *Metapopulation performance metrics: portfolio effect*

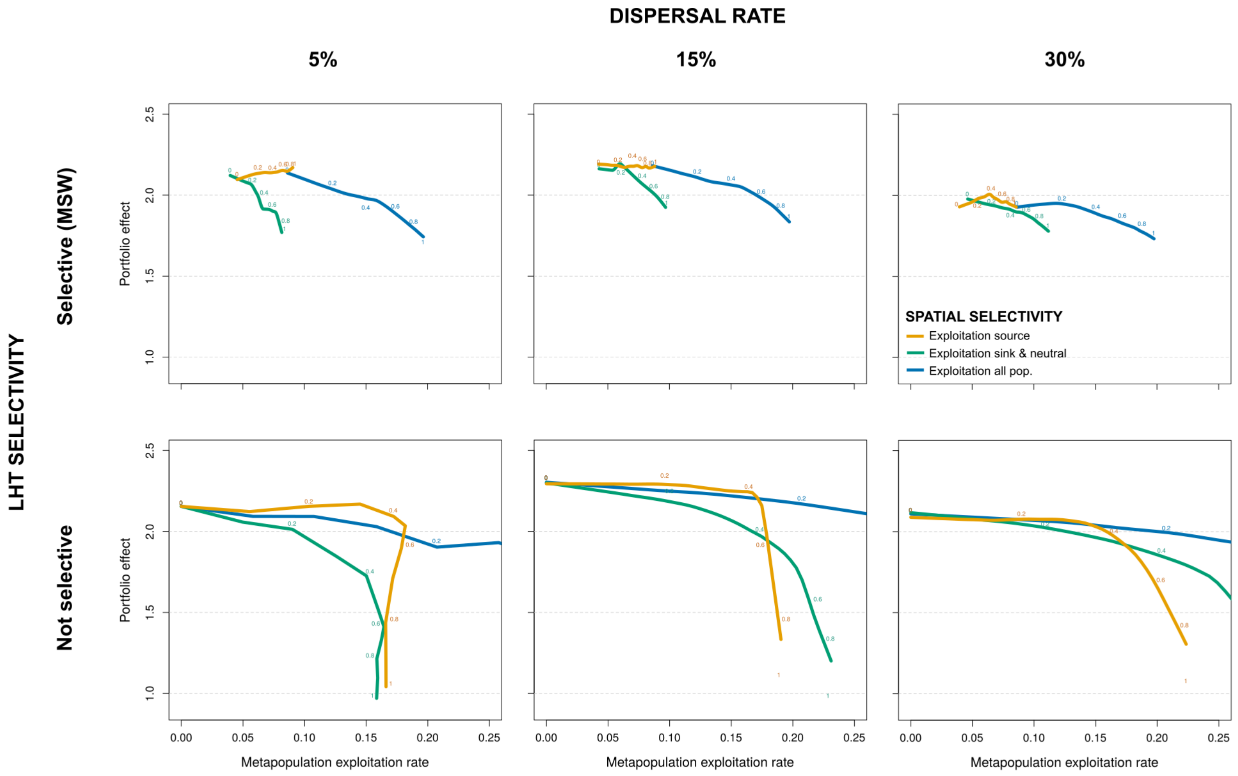


Figure S3: Metapopulation portfolio effect (over 50 years) averaged over simulations for each scenario of dispersal rate (5%, 15%, 30%), life history selectivity (top/bottom) and spatial selectivity (colors), for increasing metapopulation exploitation rate (MER). The local exploitation rates are reported adjacent to the curves (loess regression) for each scenario.

We computed the metapopulation stability using the Portfolio effect metric (hereafter $PE$) which was based on the variance of number of returns over the 50 years of simulations as in Lamarins et al. (2022*b*). The metric was then averaged over simulation replicates.

*Effect of life history selectivity.* When exploitation rate was even across populations, the decrease of portfolio effect strength with increasing exploitation was stronger in the case of selective exploitation on MSW compared to non-selective exploitation regardless of dispersal rate.

*Effect of spatial selectivity.* The scenario of even exploitation rate across all populations resulted in the highest levels of portfolio effect strength except for non selective and low dispersal. Then, the strength of portfolio effect was higher when exploitation was focused on source populations compared to focused on sink and neutral populations, except for high exploitation rates (>20%).

#### 4.8 *Metapopulation performance metrics: extinction risk*

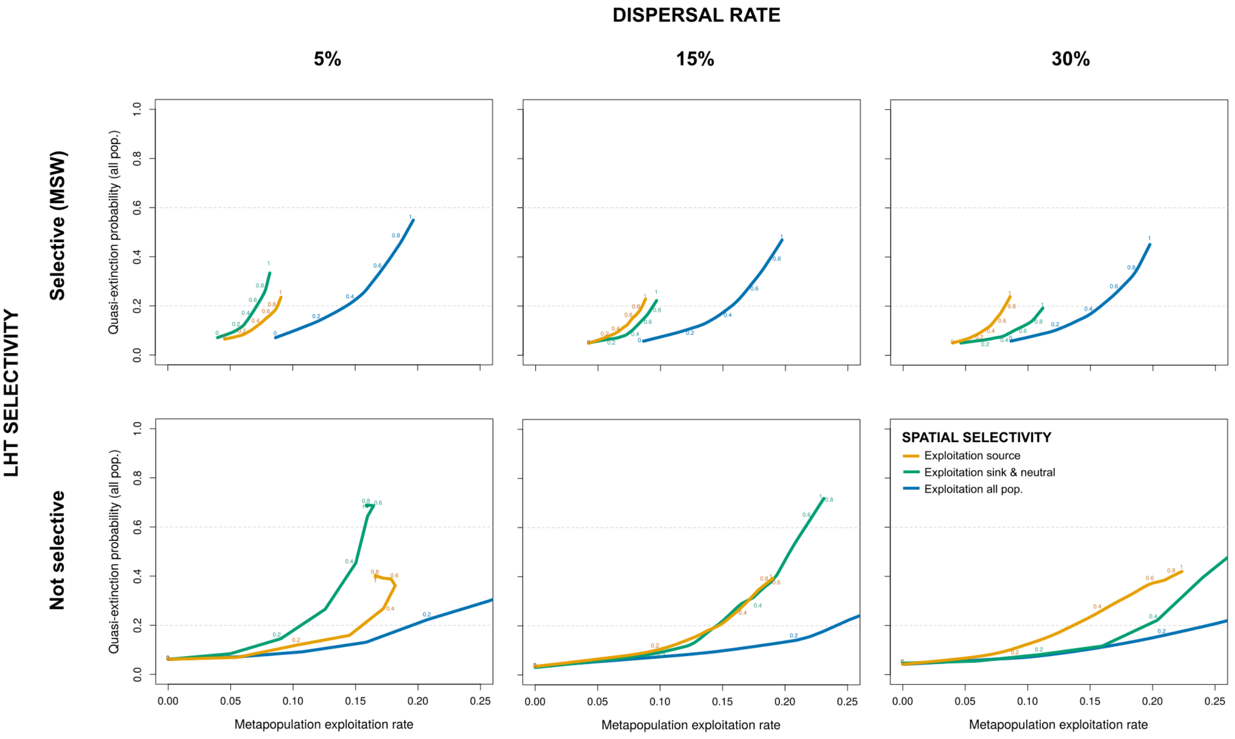


Figure S4: Quasi-extinction risk, averaged over all populations and simulations for each scenario of dispersal rate (5%, 15%, 30%), life history selectivity (top/bottom) and spatial selectivity (colors), for increasing metapopulation exploitation rate (MER). The local exploitation rates are reported adjacent to the curves (loess regression) for each scenario.

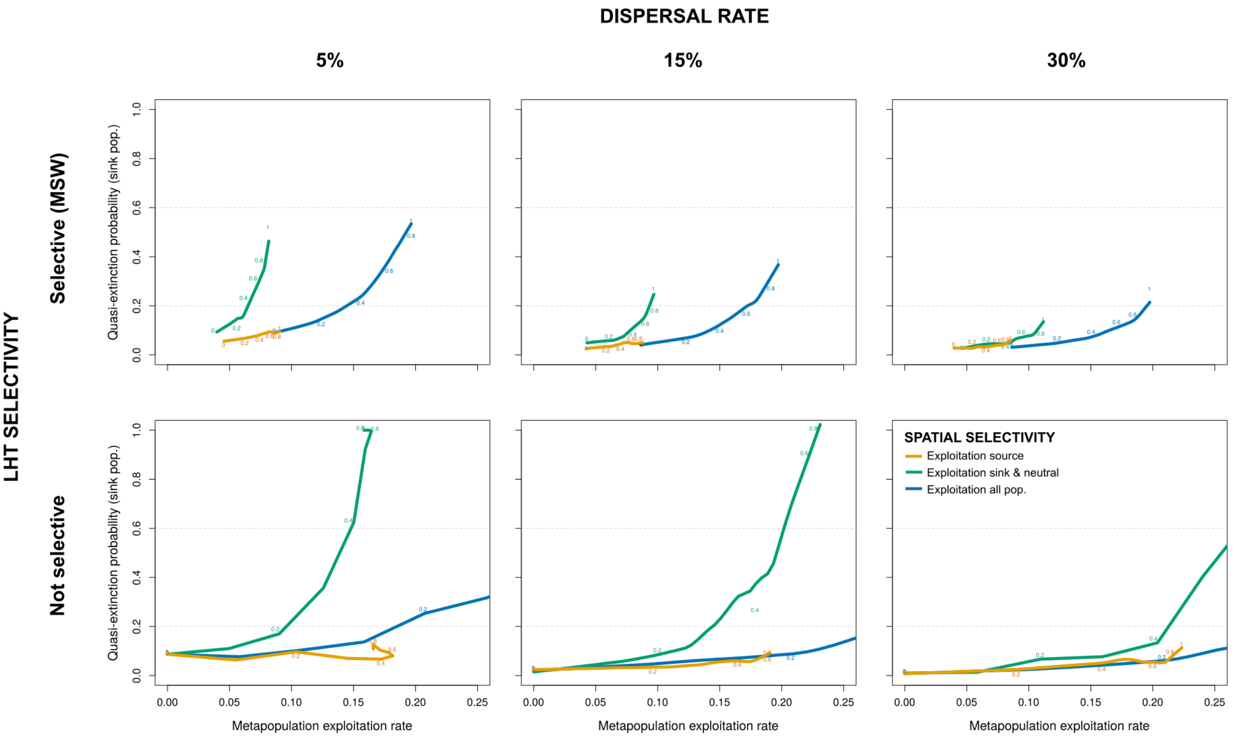


Figure S5: Quasi-extinction risk, averaged over sink populations only and simulations for each scenario of dispersal rate (5%, 15%, 30%), life history selectivity (top/bottom) and spatial selectivity (colors), for increasing metapopulation exploitation rate (MER). The local exploitation rates are reported adjacent to the curves (loess regression) for each scenario.

To evaluate the persistence of the network, we computed for each scenario the quasi-extinction risk for each population. It was computed as the proportion of simulations where the density of juveniles (parr0+) was at least 2 years consecutively below a threshold defined as 20% the population carrying capacity ($Rmax$, see Okamoto et al., 2020). We averaged the quasi-extinction risk over all populations (Fig. S4), and then over sink populations (Fig. S5) and source populations (Fig. S6) only.

*Effect of life history selectivity.* The average local extinction risk of all populations also increased with exploitation, and more strongly when exploitation was selective on MSW fish.

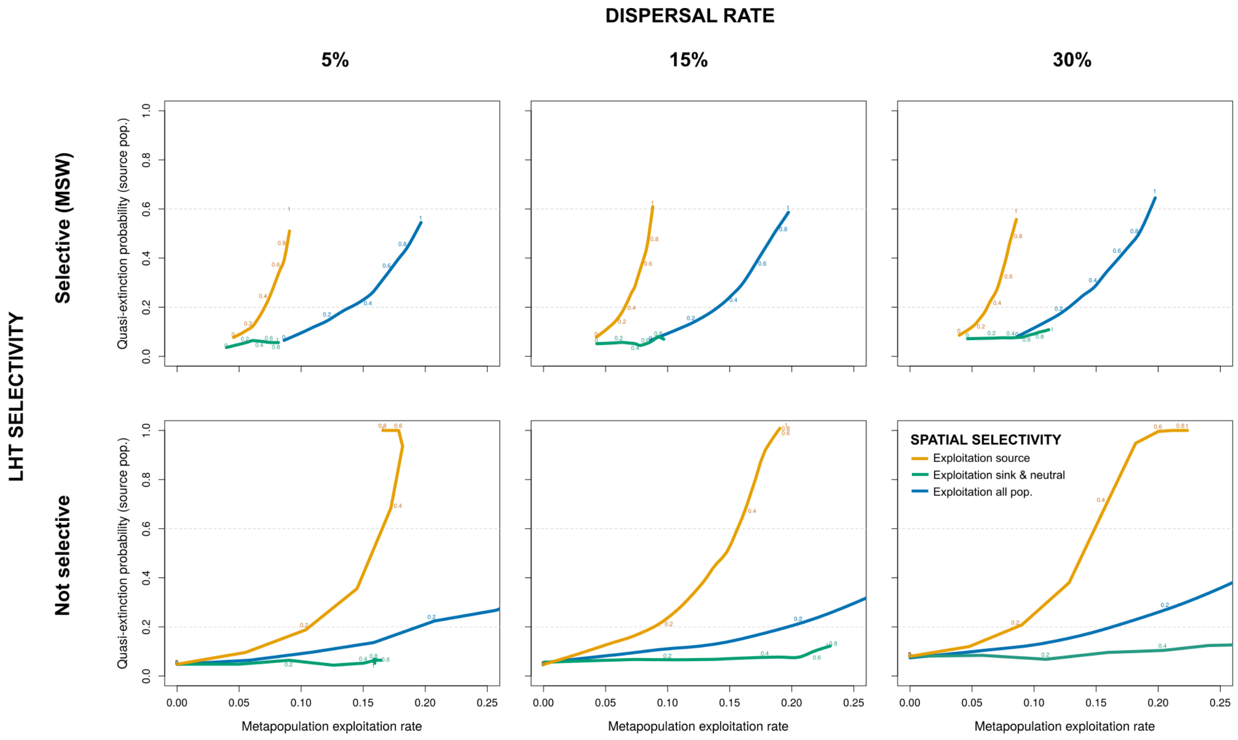


Figure S6: Quasi-extinction risk, averaged over source populations only and simulations for each scenario of dispersal rate (5%, 15%, 30%), life history selectivity (top/bottom) and spatial selectivity (colors), for increasing metapopulation exploitation rate (MER). The local exploitation rates are reported adjacent to the curves (loess regression) for each scenario.

*Effect of spatial selectivity.* The scenario of even exploitation rate across all populations resulted in the lowest levels of local population extinction risk averaged over all populations (Fig. S4), regardless of dispersal rate and scenarios of selectivity on traits. The two spatially selective strategies induced similar levels of extinction risk if dispersal rate was 15%, but there was a higher risk associated with exploitation of sink and neutral populations if dispersal was lower, whereas it was the contrary if dispersal was higher. However, by considering the extinction risk of sink populations only (Fig. S5), the best strategy was the exploitation of source populations (conservation of sink and neutral), followed by the even exploitation of all populations. Very high rates of extinction risk were reached for high exploitation rates in the case of sink and neutral exploitation (conservation of source). The opposite pattern was found by looking at the extinction risk of source populations only (Fig. S6).

#### 4.9 Exploitation metrics: total catch

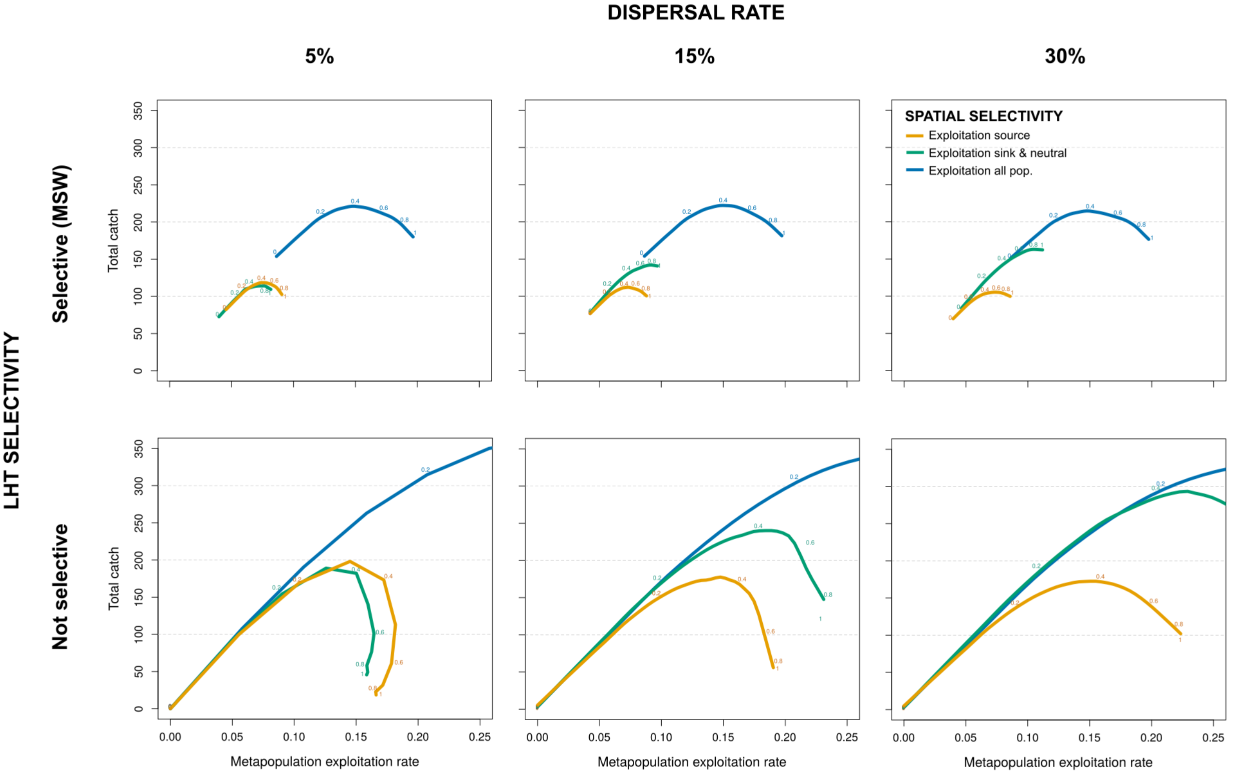


Figure S7: Metapopulation total number of catch (last 5 years) averaged over simulations for each scenario of dispersal rate (5%, 15%, 30%), life history selectivity (top/bottom) and spatial selectivity (colors), for increasing metapopulation exploitation rate (MER). The local exploitation rates are reported adjacent to the curves (loess regression) for each scenario.

Regarding exploitation, we averaged the total number of catch (i.e., summed over all populations) over the last 5 years and simulation replicates.

*Effect of life history selectivity.* Regardless of dispersal rate, exploitation selectivity on life history reduced the total number of catch especially for high exploitation rates.

*Effect of spatial selectivity.* The strategy of even exploitation rate across all populations showed the highest total catch regardless of dispersal rate and life history selectivity. Then, the strategy of exploitation of sink and neutral populations produced lower levels of total catch, but they were higher than under the source exploitation strategy for high dispersal. For lower rates of dispersal (5%), the strategy of exploitation of sink and neutral populations showed similar yield as the exploitation of source.

#### 4.10 Exploitation metrics: MSW catch

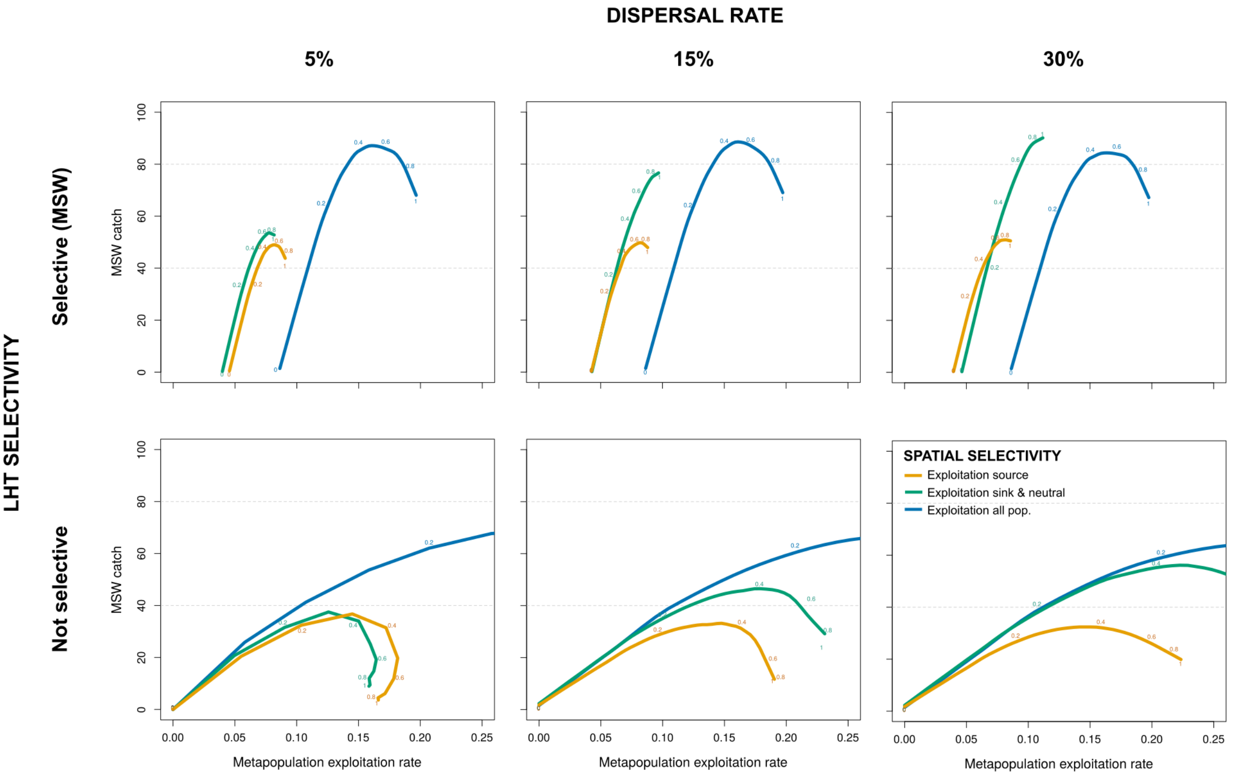


Figure S8: Metapopulation number of MSW catch (last 5 years) averaged over simulations for each scenario of dispersal rate (5%, 15%, 30%), life history selectivity (top/bottom) and spatial selectivity (colors), for increasing metapopulation exploitation rate (MER). The local exploitation rates are reported adjacent to the curves (loess regression) for each scenario.

We also averaged the total number of MSW catch (i.e., summed over all populations) over the last 5 years and simulation replicates.

*Effect of life history selectivity.* Selective exploitation on MSW led to higher number of MSW catch compared to non-selective exploitation, but it follows a decrease with increased exploitation rates after a certain threshold.

*Effect of spatial selectivity.* For non-selective exploitation on life histories, the strategy of even exploitation rate across all populations showed the highest number of MSW catch regardless of dispersal rate. Then, the strategy of exploitation of sink and neutral populations produced lower levels of catch, but they were higher than under the source exploitation strategy for high dispersal. For lower rates of dispersal (5%), the strategy of exploitation of sink and neutral populations showed similar yield as the exploitation of source. But for selective exploitation on MSW, spatially selective strategies led to a higher number of MSW catch, especially if exploitation was focused on sink and neutral populations and if dispersal rates were high.

#### *4.11 Exploitation metrics: 1SW catch*

We also averaged the total number of 1SW catch (i.e., summed over all populations) over the last 5 years and simulation replicates.

*Effect of life history selectivity.* The number of 1SW catch increased with exploitation rates if exploitation was non-selective, but it decreased if exploitation was selective on MSW, despite a constant local exploitation rate applied on 1SW (10%).

*Effect of spatial selectivity.* The strategy of even exploitation rate across all populations remained the best strategy for 1SW catch levels. Then, for low dispersal rates, similar number of 1SW catch were observed for the two spatial strategies, while for higher dispersal rates, the strategy of exploitation of sink and neutral populations showed greater number of 1SW catch than the strategy of source exploitation.

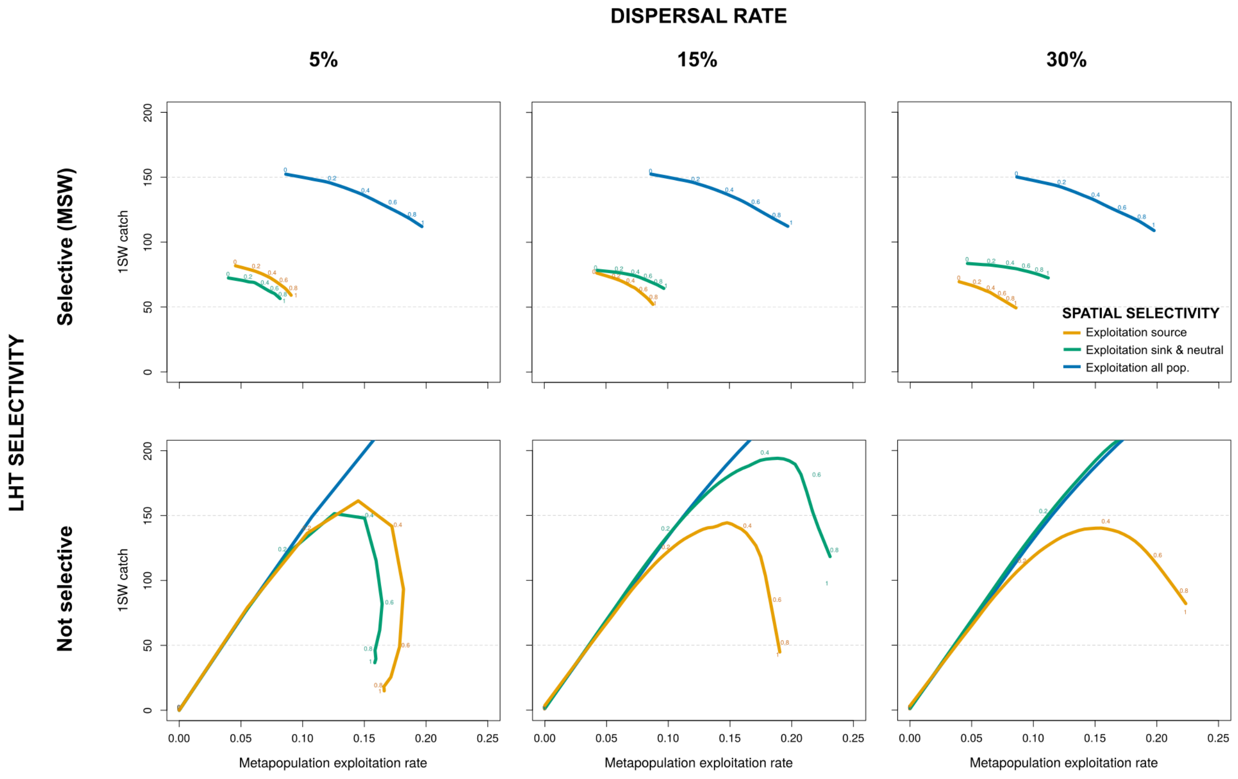


Figure S9: Metapopulation number of 1SW catch (last 5 years) averaged over simulations for each scenario of dispersal rate (5%, 15%, 30%), life history selectivity (top/bottom) and spatial selectivity (colors), for increasing metapopulation exploitation rate (MER). The local exploitation rates are reported adjacent to the curves (loess regression) for each scenario.

#### 4.12 *Evolution metrics: growth potential*

We evaluated the evolutionary dynamics of the metapopulation by looking at the average value of genetic river growth potential among philopatric individuals (i.e., excluding immigrants) over the whole metapopulation, over the last 5 years and simulation replicates.

*Effect of life history selectivity.* Increased intensity of non-selective exploitation decreased the average value of growth potential, while it increased with exploitation rates if exploitation was selective on MSW.

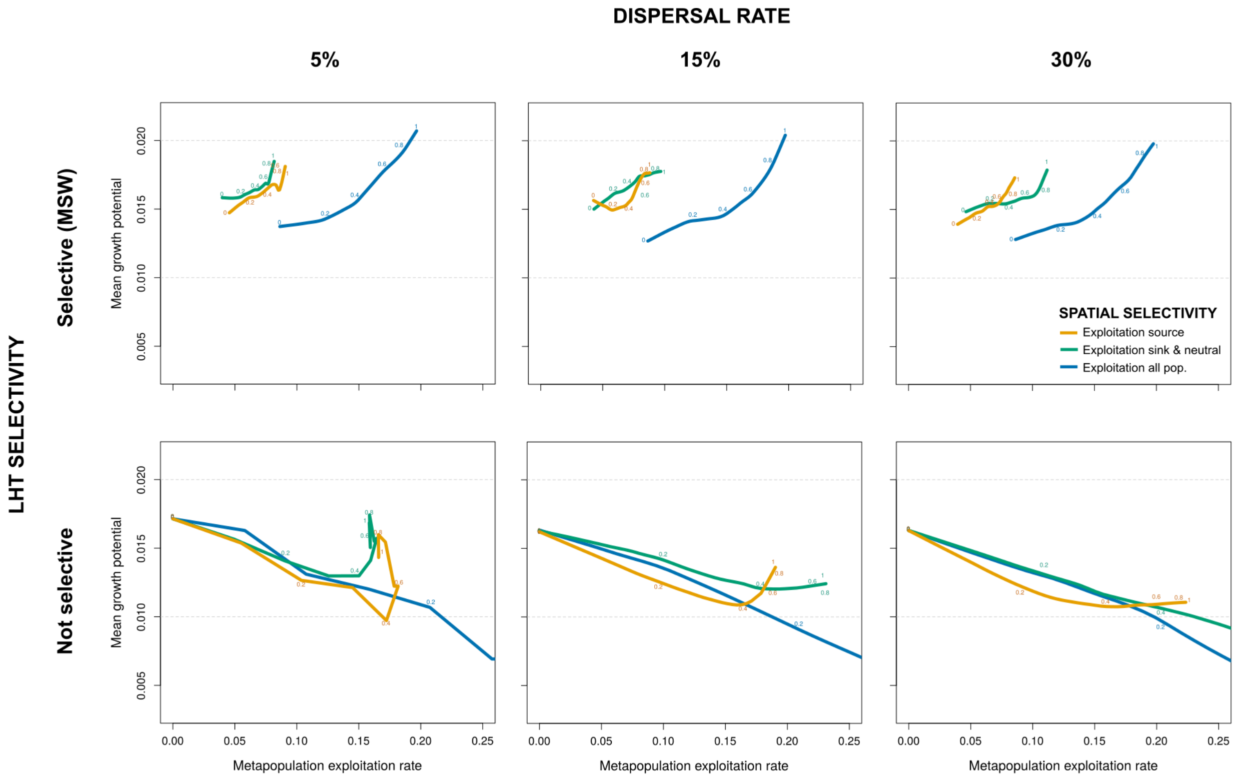


Figure S10: Average genotypic value of growth potential among philopatric returns (last 5 years) over the metapopulation and simulations for each scenario of dispersal rate (5%, 15%, 30%), life history selectivity (top/bottom) and spatial selectivity (colors), for increasing metapopulation exploitation rate (MER). The local exploitation rates are reported adjacent to the curves (loess regression) for each scenario.

*Effect of spatial selectivity.* This evolutionary response to selective exploitation on MSW appeared to be stronger if exploitation was spatially selective.

#### 4.13 Evolution metrics: maturation thresholds

We looked at the average value of female anadromous and male parr genetic maturation thresholds among philopatric individuals (i.e., excluding immigrants) over the whole metapopulation, over the last 5 years and simulation replicates.

*Effect of life history selectivity.* No evolution of female anadromous maturation threshold was observed in response to non-selective exploitation, while it decreased with increasing exploitation

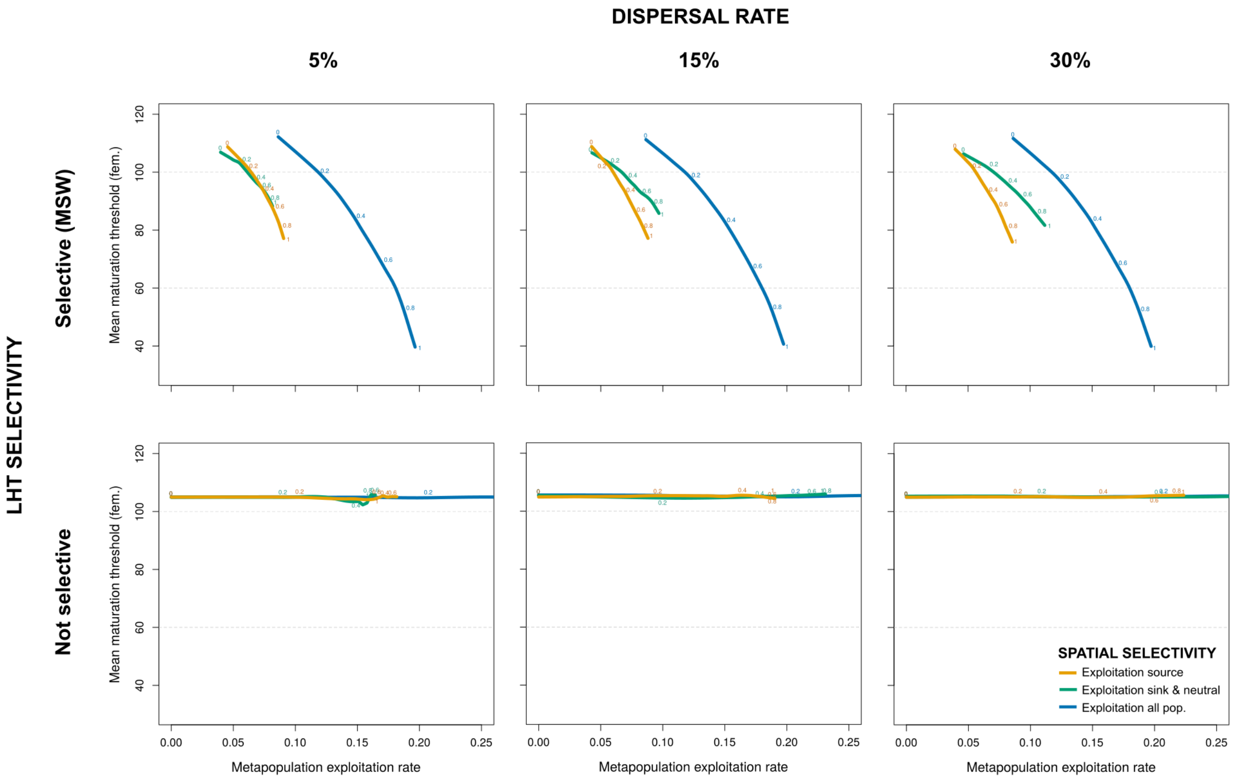


Figure S11: Average genotypic value of female anadromous maturation threshold among philopatric returns (last 5 years) over the metapopulation and simulations for each scenario of dispersal rate (5%, 15%, 30%), life history selectivity (top/bottom) and spatial selectivity (colors), for increasing metapopulation exploitation rate (MER). The local exploitation rates are reported adjacent to the curves (loess regression) for each scenario.

rates if exploitation was selective on MSW. In contrary, male parr maturation threshold decreased with increased exploitation rates when exploitation was not selective while it didn't change when exploitation was selective on MSW.

*Effect of spatial selectivity.* No effect of spatial selectivity of exploitation was found on the maturation threshold of male parr, nor on the maturation threshold of female anadromous fish in the case of non-selective exploitation. However, a stronger evolutionary response in female anadromous maturation threshold was induced by spatial exploitation when exploitation was selective on MSW, and especially when exploitation was focused on source populations with high dispersal rates.

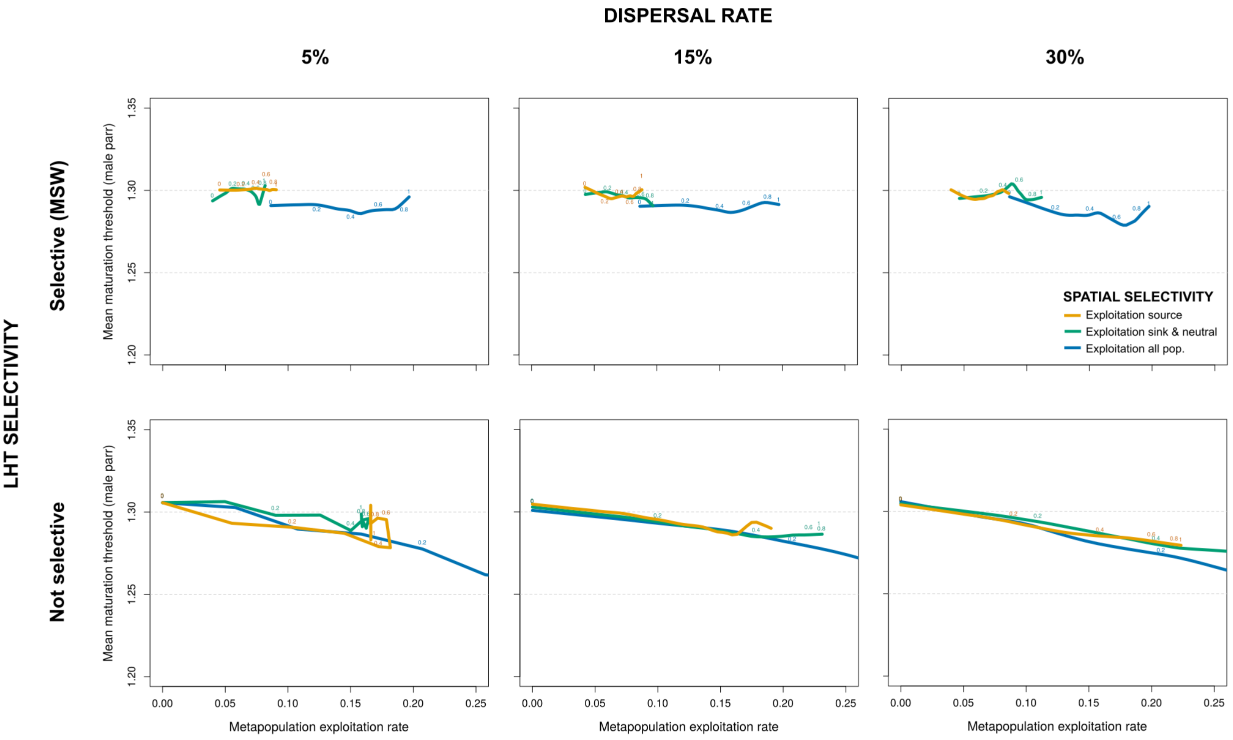


Figure S12: Average genotypic value of male parr maturation threshold among philopatric returns (last 5 years) over the metapopulation and simulations for each scenario of dispersal rate (5%, 15%, 30%), life history selectivity (top/bottom) and spatial selectivity (colors), for increasing metapopulation exploitation rate (MER). The local exploitation rates are reported adjacent to the curves (loess regression) for each scenario.

#### *4.14 Evolution metrics: ratio MSW/1SW*

For phenotypic changes, we computed the ratio between the number of MSW and 1SW returning philopatric adults (i.e., excluding immigrants) over the metapopulation, averaged over the last 5 years and simulation replicates.

*Effect of life history selectivity.* When exploitation was non-selective, there was a slight decrease of the ratio MSW/1SW with increasing exploitation rates, while it strongly decreased with exploitation rates if exploitation was selective on MSW.

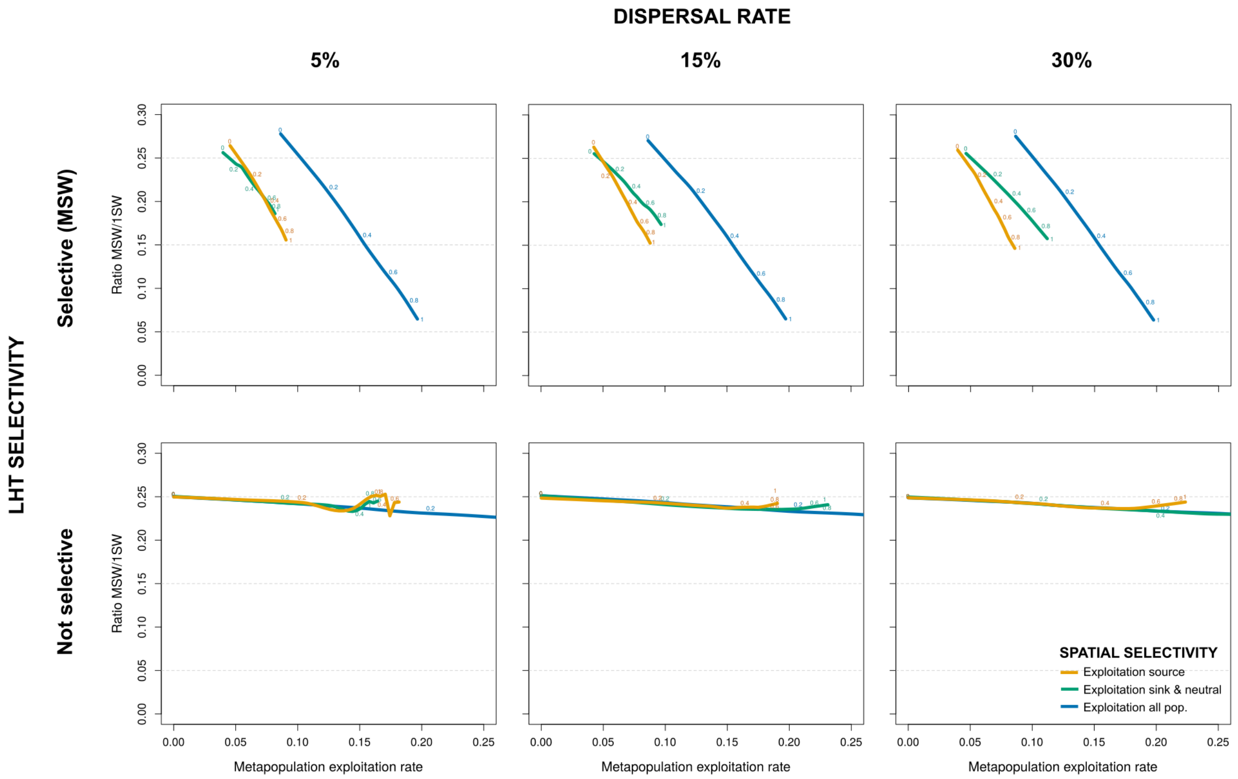


Figure S13: Ratio MSW/1SW (last 5 years) over the metapopulation and simulations for each scenario of dispersal rate (5%, 15%, 30%), life history selectivity (top/bottom) and spatial selectivity (colors), for increasing metapopulation exploitation rate (MER). The local exploitation rates are reported adjacent to the curves (loess regression) for each scenario.

*Effect of spatial selectivity.* This evolutionary response to selective exploitation on MSW was stronger if exploitation was spatially selective, especially if exploitation was focused on source populations with high dispersal rates.

#### 4.15 Evolution metrics: genetic variability

We also looked at the variation in genetic traits within the metapopulation by calculating the standard deviation of genetic river growth potential and female anadromous maturation threshold among philopatric returns (i.e., excluding immigrants), which was averaged over the last 5 years and simulation replicates.

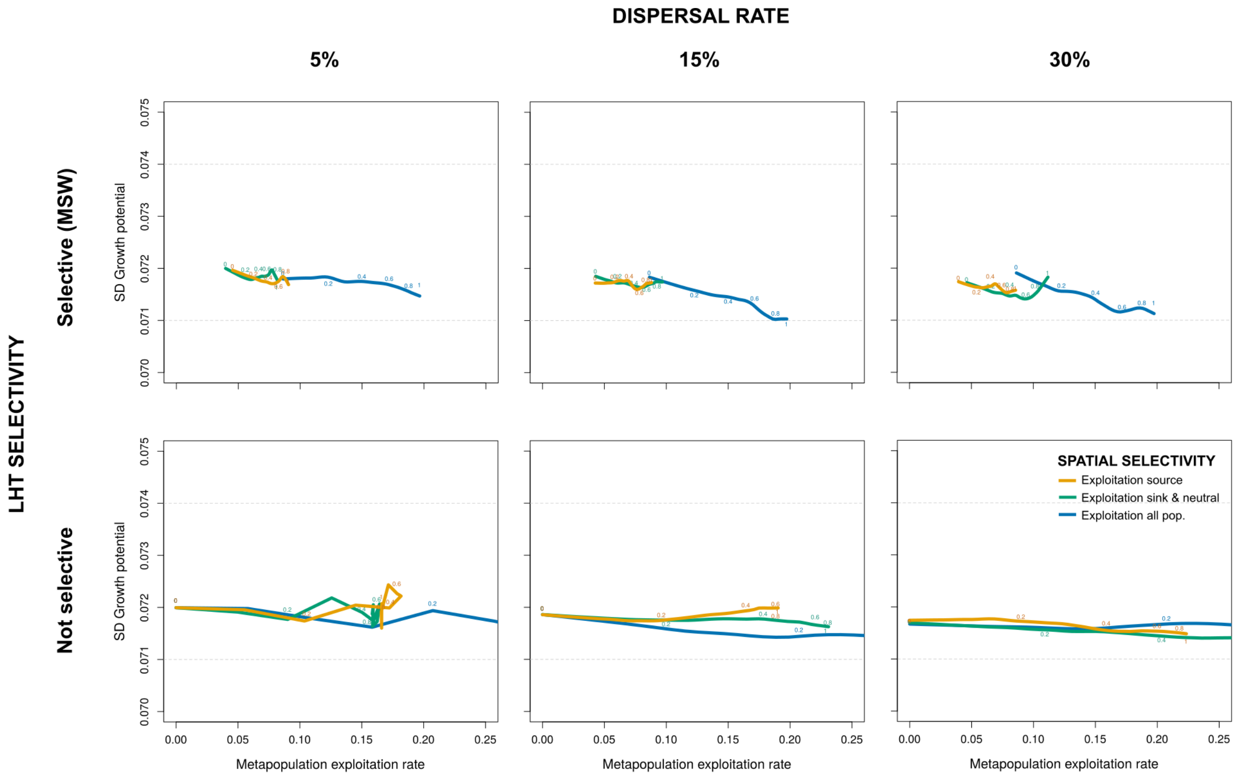


Figure S14: Standard deviation of the genotypic value of growth potential among philopatric returns (last 5 years) over the metapopulation and simulations for each scenario of dispersal rate (5%, 15%, 30%), life history selectivity (top/bottom) and spatial selectivity (colors), for increasing metapopulation exploitation rate (MER). The local exploitation rates are reported adjacent to the curves (loess regression) for each scenario.

*Effect of life history selectivity.* For growth potential, there was no clear difference in the genetic variability over the metapopulation between scenarios of exploitation selectivity on traits. However, there was a decrease of genetic variability of maturation threshold with increased exploitation rates when exploitation was selective on MSW, and this was the case regardless dispersal rate.

*Effect of spatial selectivity.* For growth potential, there was no clear difference in the genetic variability over the metapopulation between scenarios of exploitation spatial selectivity. However, when exploitation was selective on MSW, spatial selectivity increased the overall genetic variability of maturation threshold, especially for low dispersal rates.

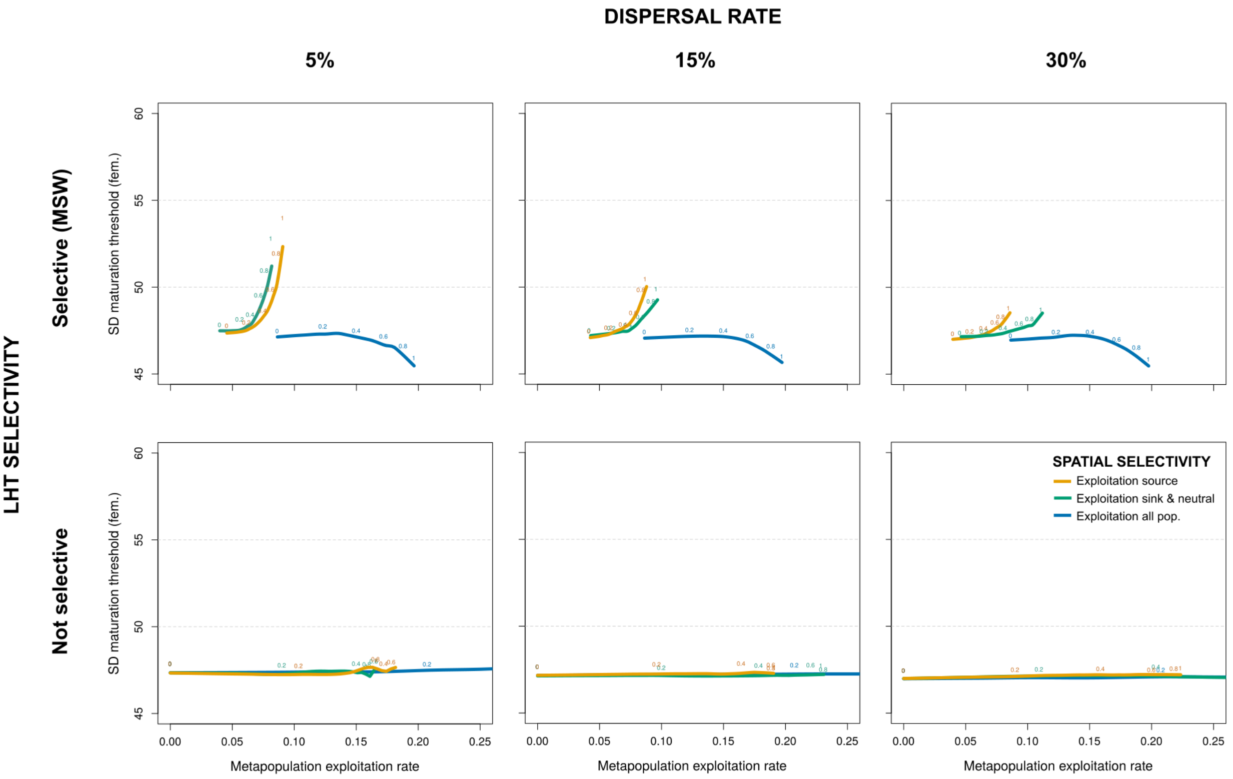


Figure S15: Standard deviation of the genotypic value of female anadromous maturation threshold among philopatric returns (last 5 years) over the metapopulation and simulations for each scenario of dispersal rate (5%, 15%, 30%), life history selectivity (top/bottom) and spatial selectivity (colors), for increasing metapopulation exploitation rate (MER). The local exploitation rates are reported adjacent to the curves (loess regression) for each scenario.
